# Artificial light at night induces stress and affects evolutionary change in bats

**DOI:** 10.64898/2025.12.28.696742

**Authors:** Tom L. Jenkins, Thomas Foxley, Yury Malovichko, Emma L. Stone, Orly Razgour

## Abstract

Artificial light at night (ALAN), a pervasive component of urbanization, is rapidly transforming nocturnal environments and imposing novel selection pressures on wildlife. While behavioral responses to ALAN are increasingly well documented, physiological and evolutionary consequences are not well understood, limiting species conservation management. We used a multi-omics framework to investigate the ecological and evolutionary consequences of ALAN on a light sensitive species, the lesser horseshoe bat (*Rhinolophus hipposideros*), combining experiments with transcriptomics, population genomics and landscape ecology. Experimental light exposure resulted in 252 differentially expressed genes, enriched for functions in DNA damage repair, apoptosis regulation and oxidative stress mitigation. Whole genome sequencing identified genetic variation associated with ALAN and urbanization, including variants in genes linked to visual and neural function. Landscape analysis revealed that ALAN and distance to broadleaf woodland best explained patterns of population connectivity. These findings demonstrate the possible drivers behind behavioral responses, showing that ALAN can act both as an acute molecular stressor and contribute to evolutionary change, with possible consequences for movement ecology and local adaptation. Given the accelerating expansion of urbanization, understanding species-specific responses to ALAN across molecular, demographic and evolutionary timescales is critical for predicting outcomes for biodiversity and informing urban planning.

## Introduction

Artificial light at night (ALAN), including streetlights, vehicle headlights and commercial and residential illumination, is a hallmark of urbanization, transforming nightscapes across the globe. While ALAN plays important roles in modern human society, such as enabling activity at night and improving safety, it also represents a pervasive environmental pollutant(1), with negative effects reported for ecological and human health(2). Light pollution has been increasing rapidly, both in intensity and spatial coverage, with a 10 % annual increase in average night sky brightness between 2011 and 2022(3). ALAN has profound implications for biodiversity, particularly for nocturnal animals, whose biology and behavior are closely linked to natural light-dark cycles(4). ALAN disrupts circadian rhythms(5), disturbs nocturnal pollination networks(6), and can affect patterns of foraging, reproduction(7, 8), migration(9), and predator-prey interactions(10). Since nocturnal species evolved under predictable cycles of daylight and darkness, ALAN introduces a novel environmental stressor, one that many species are not physiologically or behaviorally adapted to deal with.

Bats, as highly mobile, nocturnal flying mammals, represent a key taxonomic group for understanding the biological consequences of ALAN. They are essential to ecosystem functioning through their roles as pollinators, seed dispersers and insect predators(11). Bats have behavioral adaptations to darkness, such as echolocation, low-light vision, and roosting strategies that help them avoid diurnal predators. However, ALAN has been shown to affect these natural behaviors(12). Some species, particularly light-averse species, such as horseshoe bats (*Rhinolophus* sp.), may delay emergence, reduce foraging activity, or abandon roosts entirely(12, 13). Disturbance from ALAN can compromise energy intake, reduce reproductive success, and ultimately affect population viability. Conversely, bat species that appear less sensitive to ALAN, due to their fast and agile flight ability, may benefit from lit environments where flying insects are concentrated around artificial lights(14), though even these ‘light tolerant’ species will avoid light when possible(15). This highlights the complex and species-specific responses to ALAN.

While behavioral and observational studies have provided insight into how bats and other nocturnal animals respond to ALAN, we are yet to understand the impact of ALAN on underlying physiological mechanisms, particularly at the molecular level (but see(13)), and how these may affect bat fitness and survival. In many urban environments, bats may tolerate ALAN where it is unavoidable and may experience more subtle molecular and physiological stress responses, which could have a knock-on effect on long-term survival and reproduction. Investigating the underlying drivers of behavioral responses to ALAN is the logical next step to understand whether ALAN results in fitness costs which may have population level impacts that may threaten bat conservation status. In addition, no study has investigated whether bat populations have evolved adaptations to increasing levels of ALAN, the discovery of which would demonstrate that ALAN influences the evolutionary trajectory of species. These knowledge gaps have, in part, been impeded by the lack of genomic resources for bat species. However, recent initiatives, such as the Bat1K project(16), are generating and releasing annotated chromosome-level reference genomes for bats.

In this study we use a multi-omics approach (transcriptomics and genomics) to assess the impact of ALAN on gene expression, gene flow and adaptation in the lesser horseshoe bat (*Rhinolophus hipposideros*), a threatened species in the UK known to avoid ALAN along its commuting routes(17). We experimentally tested if bats exposed to ALAN on emergence from maternity roosts showed differential gene expression compared to bats emerging in naturally dark conditions (Figure 1) to provide insights into molecular stress responses caused by ALAN and highlight the physiological consequences. For assessing long-term evolutionary changes, we sequenced whole genomes of bats from populations with varying levels of ALAN surrounding roosts to: (i) explore the influence of ALAN on gene flow and functional connectivity, and (ii) find genetic variants that are linked to genes with known functions in mammalian vision or the nervous system.

**Figure 1.**
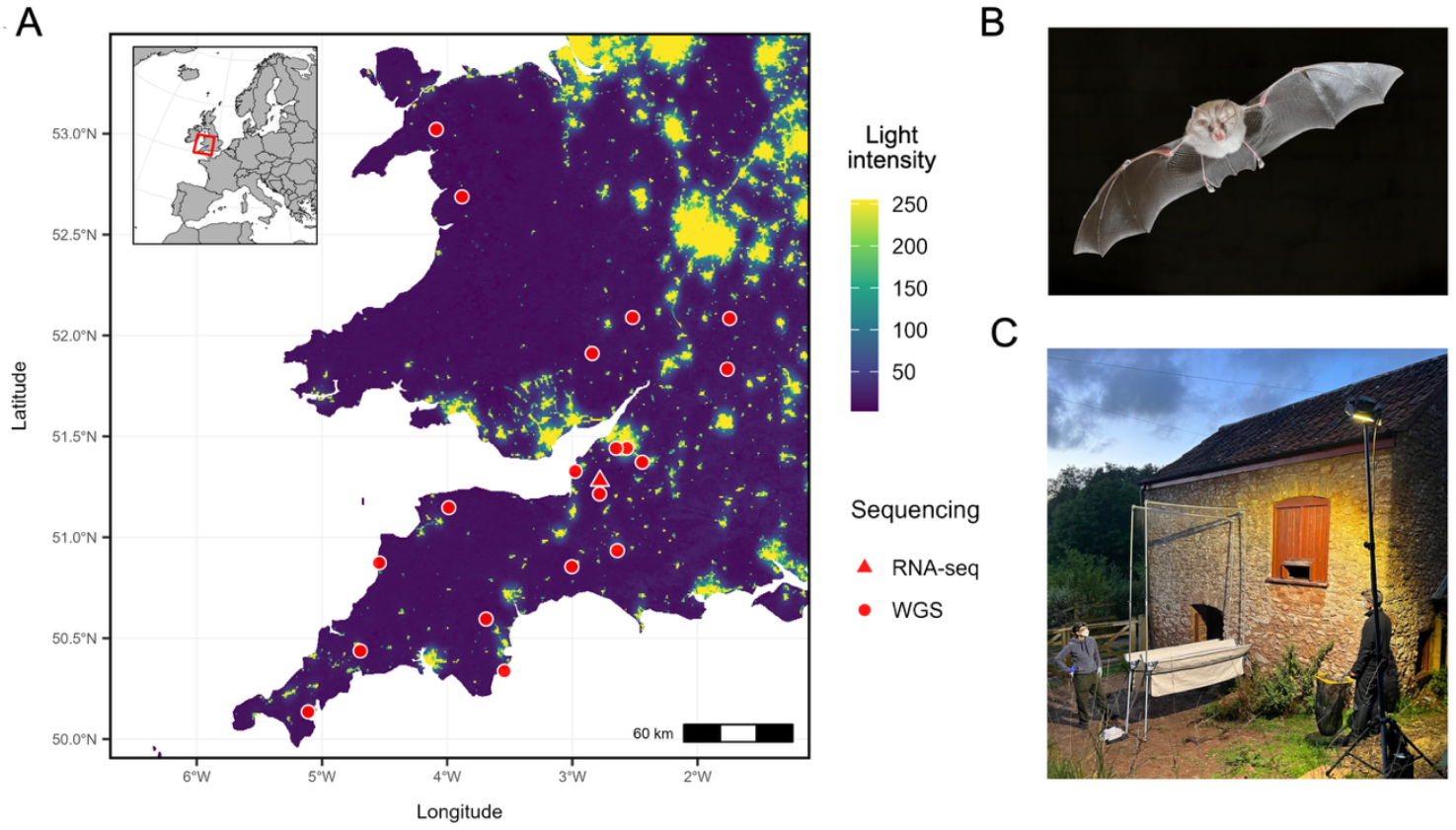
Distribution of artificial light at night (ALAN) and the sampling design. (A) Map of sampling locations in England and Wales. The single triangular point represents the gene expression experiment field site (RNA-seq) in Somerset, while the remaining circular points represent sites sampled for whole genome sequencing (WGS). The background shading represents light intensity from lower levels (dark blues) to higher levels (greens and yellows). (B) Image of a lesser horseshoe bat, Rhinolophus hipposideros (image credit: Daniel Whitby). (C) Photo of the experimental field site and the light experiment set up (photograph of the authors).

## Results

### Differential gene expression in light-exposed bats

RNA sequencing of 10 light exposed versus 10 control (natural dark conditions) *R. hipposideros* bats resulted in a median of approximately 70 million filtered reads per sample (individual bat). After removing transcripts with fewer than ten counts over all samples, transcript count data remained for 13,960 genes (out of 16,519 genes annotated in the *R. hipposideros* genome). The single factor gene expression model identified 252 differentially expressed genes (DEGs) between light and dark conditions (Table S1). All DEGs were upregulated in the light-exposed bats except for a single gene that was downregulated. For the 251 upregulated genes, while all of the control bats in dark conditions consistently showed little or no gene expression changes in the experiment, the light-exposed bats showed variability in their levels of upregulation (Figure 2). For example, relative to the mean expression level of control bats, 4-6 light-exposed bats (CH339, CH235, CH328, CH308, CH150, CH315) were upregulated in 116 genes, while the remaining light-exposed bats showed little differences in expression compared to the control bats. These patterns of biological variability are reflected in a principal components analysis (PCA) of the normalized counts for DEGs (Figure S1). This PCA shows no distinct clustering of light and dark treatments, but PC1 separates three light-exposed samples (CH339, CH235, CH328) and PC2 separates a single light-exposed sample (CH315), while all of the control samples are generally clustered together.

**Figure 2.**
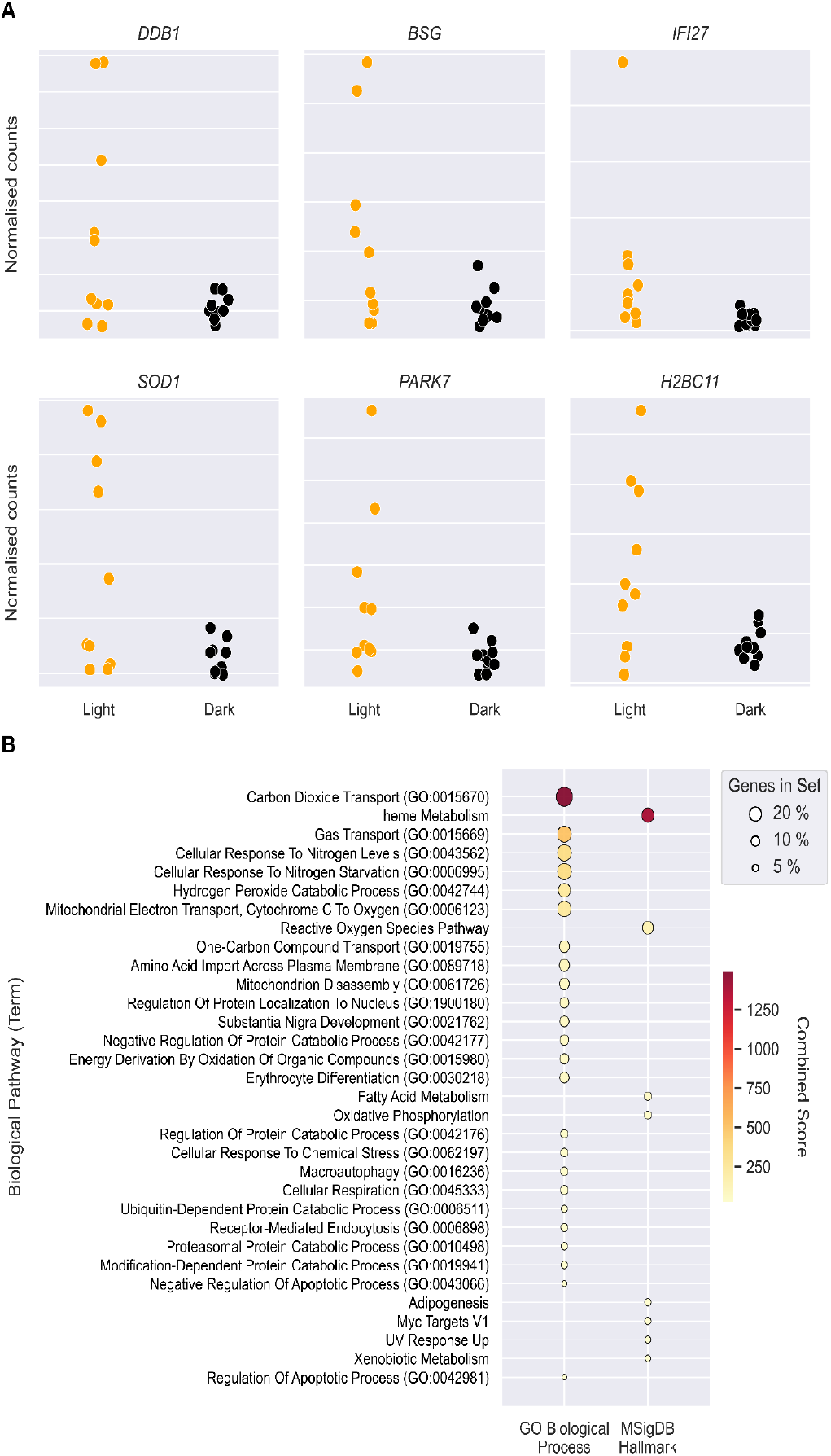
Results of the differential gene expression analysis of bats experimentally exposed to ALAN. (A) Normalized counts for a subset of upregulated genes in light-exposed bats. Each point represents the counts for an individual bat. Colors denote whether the bat was exposed to artificial lighting (orange) or was not exposed (black). (B) Biological pathways that are statistically over-represented among the differentially expressed genes are shown. Pathways are grouped by two gene set libraries: the Gene Ontology (GO) Biological Process library, which organizes genes by the biological processes, and the Molecular Signatures Database (MSigDB) Hallmark library, which summarizes well-defined biological states and processes.

Several of the upregulated DEGs are known to have important direct or indirect roles in: (i) recognising and initiating repair of UV-induced DNA damage (*DDB1, GADD45A*)(18, 19), (ii) survival of retinal cone photoreceptors (*BSG*)(20), (iii) maintenance and DNA repair (*H2BC11, MACROD1, MACROH2A1, UBE2B*)(21), (iv) apoptosis regulation (*C1D, IFI27*)(22), (v) detoxification of superoxide radicals (*CAT, SOD1*)(23), and (vi) counteracting damage from oxidative stress (*PARK7, PDK2, PRDX2, SIRT2*)(24). The biological pathway enrichment analysis identified 36 over-represented pathways (Table S2). These pathways reflected many of the functional roles of the upregulated genes outlined above, such as elevated response to UV (UV Response Up), regulation of apoptosis (GO:0043066, GO:0042981), reactive oxygen species (ROS) and the breakdown of hydrogen peroxide (Reactive Oxygen Species Pathway, GO:0000302, GO:0042744), and responses to oxidative stress (GO:0034599, GO:0062197) (Figure 2).

### Population structure and genomic diversity

Whole genome sequencing of 198 bats resulted in 81,169 high-quality (<10% missing genotypes) biallelic single nucleotide polymorphisms (SNPs), of which 81,104 were identified as putatively neutral SNPs and used in the analysis of population structure and landscape genetics. A PCA of the SNP genotypes showed a spatial genetic cline from south-west Cornwall to south-east Wales and the south-western border with England (Figure 3A). It also showed distinct separation of two sites in North Wales (NW1 & NW2) compared to the main cluster gradient of sites. These results were supported by admixture analysis, which identified three ancestral populations (*K*3) based on the cross-entropy criterion (Figure S2), and similar clinal patterns of genetic clustering (Figure 3B).

**Figure 3.**
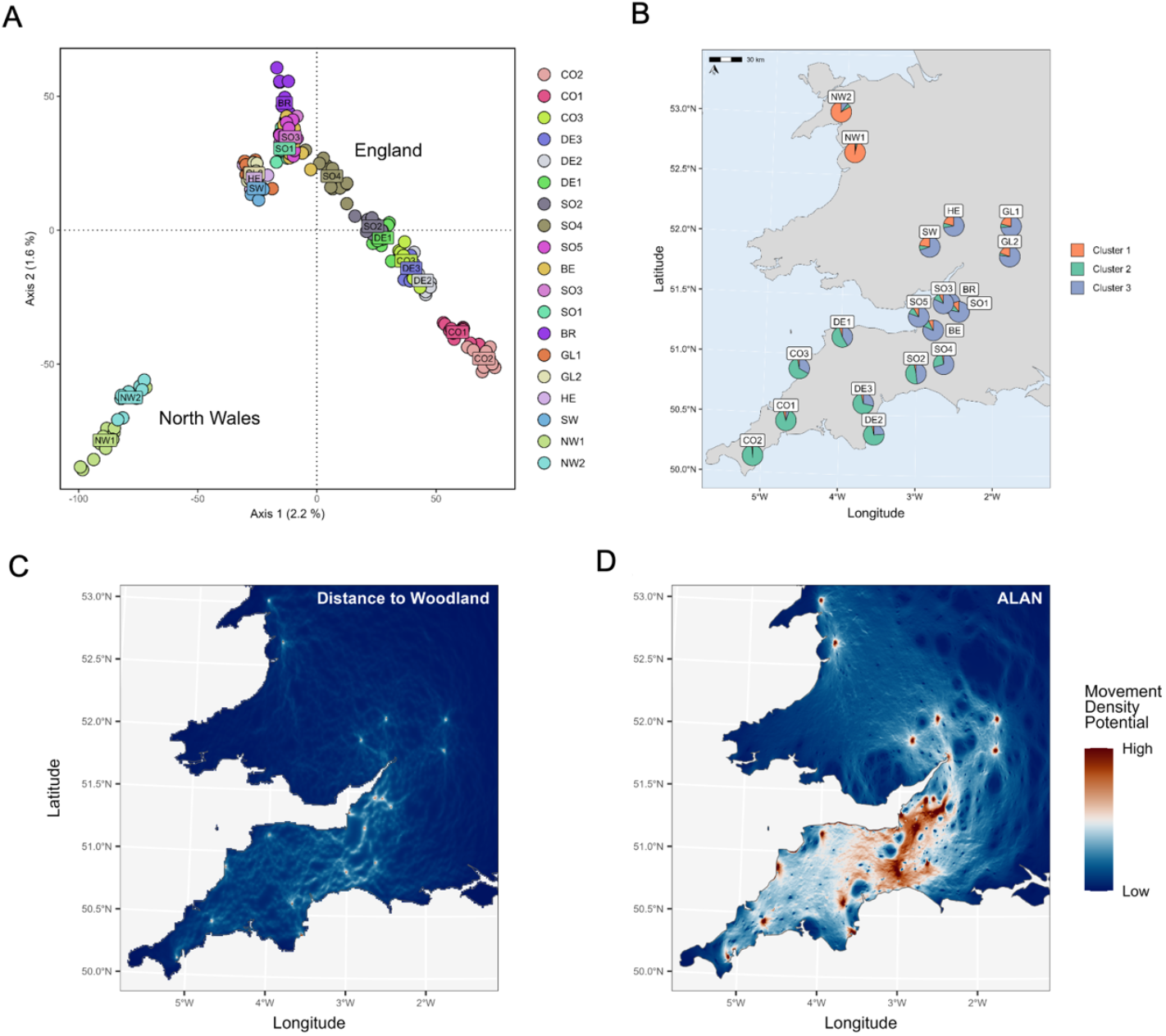
Genetic population structure of *R. hipposideros* across Britain. (A) Principal components analysis on the genotypes of 81,104 putatively neutral single nucleotide polymorphisms (SNPs). Colored points represent the site where bats were sampled (See Table S2 for site information). (B) Results of the population admixture analysis at the site (roost) level, showing average assignment of individual bats in each site to the three genetic clusters based on their proportion of ancestry in each cluster. (C-D) Results of the landscape genetics analysis identifying barriers to genetic connectivity between *R. hipposideros* colonies across Britain, showing predicted movement density potential between colonies based on the effect of landscape resistance due to distance to woodland (C) and the intensity of ALAN (D). In (C) and (D), maps show electric currents crossing the matrix between pairs of colonies and current density may be interpreted as movement density potential.

Genome-wide heterozygosity (theta), calculated by examining base positions in the genome alignments and excluding runs of homozygosity(25), ranged from 0.0017–0.0026 over all individuals. When averaged across sites, lower values of theta were generally associated with sites at the edge of the sampling distribution (e.g. North Wales and south-west Cornwall) compared to sites at the center of the species’ geographic range (Table S3). Observed heterozygosity using the neutral SNP dataset ranged from 0.075–0.173 and average values across sites showed a similar center-edge spatial pattern as theta. There was no spatial pattern observed with the proportion or lengths of runs of homozygosity, an indicator of levels of inbreeding (Table S4). In addition, we found no significant correlations (*r* ^2^±0.30, *p*>0.05) between any metric of genomic diversity and ALAN or other landscape variables (Table S4).

### Effects of ALAN and landscape on genetic differentiation

We tested for isolation-by-distance (IBD) and found a high correlation between pairwise genetic differentiation (*F*_ST_) and Euclidean distances (*r* ^2^=0.52, *p*<0.001). To account for the effects of IBD when analyzing the effects of landscape variables on genetic connectivity, *F*_ST_ was divided by the Euclidean distance between sites in the landscape genetics analysis. After controlling for multicollinearity, the following landscape variables were compared with maximum likelihood population mixed-effects models: (i) ALAN, (ii) distance to broadleaf woodland, and (iii) a land cover scenario where woodland and grasslands do not form strong barriers but other habitat classes do, with urban areas being the strongest barrier (Table S5). The best supported landscape model was the combined effect of ALAN and distance to woodland (*r* ^2^=0.33, AICc weights=0.738, BIC weights=0.907) (Table S6). The model with only distance to woodland explained a much larger amount of the variation (*r* ^2^=0.31) than the model with ALAN alone (*r*^2^=0.02). For both variables, predicted movement density was highest around the core of the species’ range, across Somerset and Devon, and lower between sites at the edges of the range, especially between North Wales and the rest of the sites (Figure 3C-D).

### Adaptive variation in genes with functional roles in the eye and nervous system

A genotype-environment association (GEA) analysis was carried out using ALAN and urbanization as the predictor variables. Out of 81,169 SNPs, 1,554 SNPs (1.91%) were identified as associated with the predictor variables, and therefore putatively adaptive. These SNPs were distributed over all 28 autosomal chromosomes and the X chromosome. We identified 577 putatively adaptive SNPs that were located directly in or within 1kb of a gene annotation in the *R. hipposideros* genome (Figure 4; Table S7). These SNPs were located in 448 unique genes. A biological pathway enrichment analysis identified 13 over-represented pathways (Table S8), including Retinal Ganglion Cell Axon Guidance (GO:0031290) and a number of signaling, synapse regulation and sensory pathways (Figure 4).

**Figure 4.**
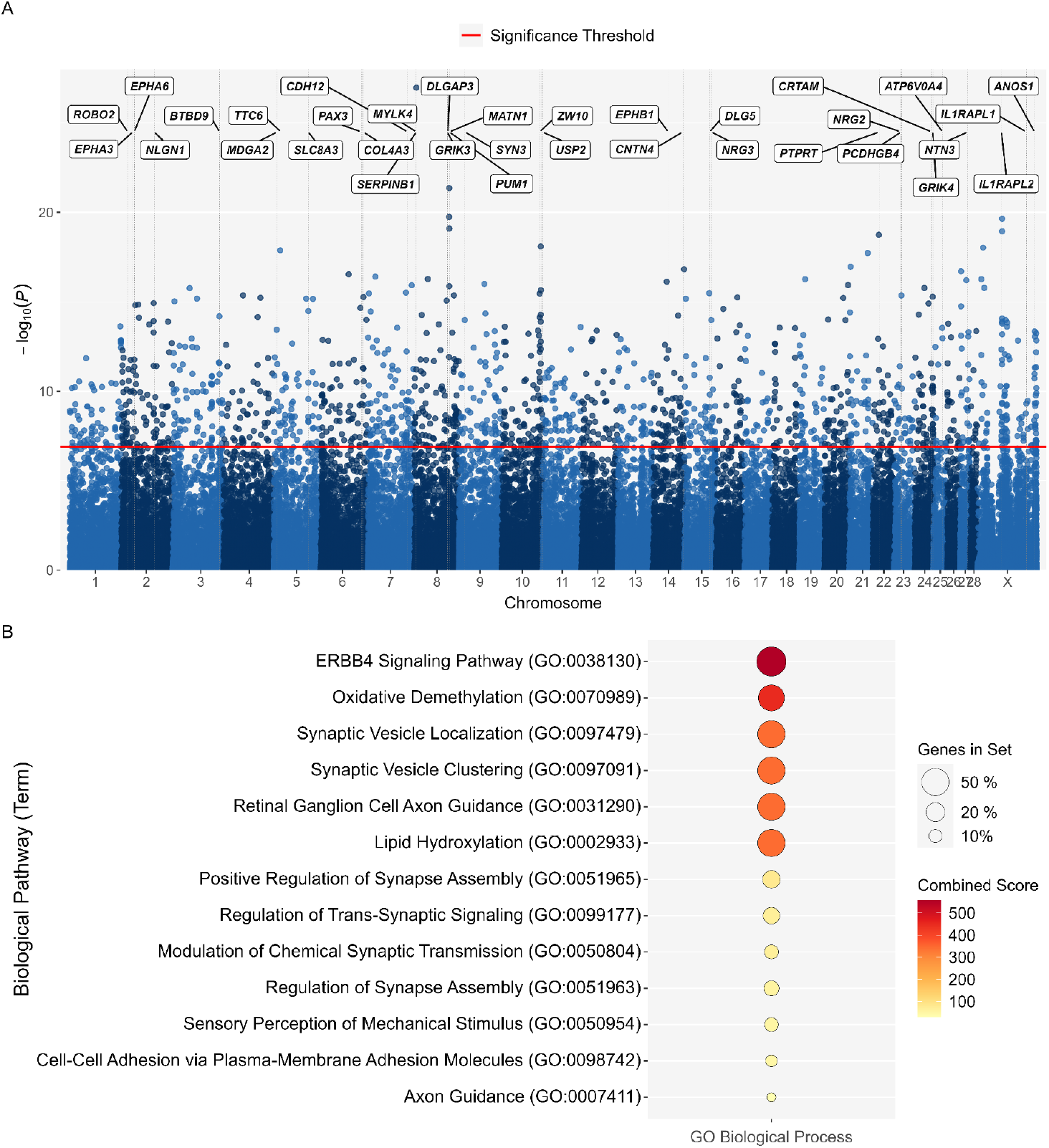
Genotype-environment association (GEA) analysis using ALAN and urban cover as the predictor variables. (A) Manhattan plot where each point represents adjusted P-values (-log10 transformed) for each single nucleotide polymorphism (SNP) on each chromosome. The red horizontal line is the threshold used for defining outlier SNPs that significantly correlate with artificial lighting and urban cover in the GEA analysis. The position of a subset of genes containing outlier SNPs (or within 1kp) are highlighted. (B) Biological pathways that are statistically over-represented among the genes containing outlier SNPs. Only enriched pathways for the Gene Ontology (GO) Biological Process library are shown as no pathways for the Molecular Signatures Database (MSigDB) Hallmark library were enriched in this analysis.

## Discussion

Artificial light at night (ALAN) is an increasingly prevalent feature of urbanized landscapes, modifying natural light regimes that have remained stable over evolutionary timescales. Here we provide evidence that even short exposure to ALAN triggers changes in gene expression in a threatened bat species, known to avoid ALAN, with potential negative consequences for fitness and survival. We found 252 genes that were differentially expressed, all but one were upregulated in light-exposed bats, indicating an acute molecular stress response involving DNA damage repair, apoptosis regulation and oxidative stress mitigation. These functional signatures are consistent with the expected physiological challenges posed by ALAN, such as increased retinal stress and elevated metabolic demands associated with altered nocturnal behavior. While such responses have been identified in other animals exposed to ALAN under laboratory or controlled conditions(7, 26), we provide the first evidence using transcriptomics in a wild ecologically realistic setting.

The variability in expression observed among light-exposed bats, with only a subset of individuals exhibiting strong responses, suggests either individual-level differences in sensitivity or an influence of unmeasured environmental or physiological variables at the time of sampling. This heterogeneity may reflect the high biological variance typical of gene expression studies of wild populations and natural (field) conditions(27). For example, it may reflect prior exposure to ALAN at the field site or small differences in exposure to light during emergence and capture. Re-exposing individuals to artificial light could reveal whether these patterns are consistent among individuals and if such molecular responses accumulate over time. Experiments with repeated exposure to light may elucidate the role of epigenetics in acclimation, which is increasingly recognized as playing a role in mediating stress responses(28, 29). Moreover, sampling eye and brain tissue, which was not possible in our study due to ethical and conservation reasons, may tell us whether the observed shifts in gene expression originate in sensory systems directly detecting light, or in systemic stress pathways.

In addition to our findings indicating strong stress responses in the short-term, our genomic analyses allowed us to extend our understanding of the impacts of ALAN on long-term evolutionary processes. We identify genomic variants associated with ALAN and urbanization, some of which were located in genes with known functional roles in the mammalian eye and nervous system. This suggests that ALAN may be shaping allele frequency variation consistent with local adaptation or selection pressures. This finding aligns with broader evidence from other animals where, for example, ALAN has been linked to evolutionary changes in circadian clock genes(5, 26). However, because the gene annotations for the *R. hipposideros* reference genome are extrapolated from model species (e.g. humans, mouse, etc.), as is commonly the practice with non-model organisms, annotated genes may not always match the functions documented for the model species. Thus, experimental validation, such as confirming changes in protein structure or knock-out studies demonstrating causal roles in ALAN response, is needed to ascertain whether these genomic variants translate to functional differences in physiology or light perception.

Isolation-by-distance, driven by the restricted dispersal ability of the species(30), is the most likely explanation for the genetic cline observed in the population structure of *R. hipposideros*. Genetic connectivity in Somerset, at the core of the species’ range, can be explained by proximity to caves (e.g. the Mendips), where bats gather to mate during autumn swarming(31) and hibernate over winter. Reduced genome-wide diversity and stronger population structure at the northern range edge, may represent relatively recent range expansion to North Wales and resulting loss of genetic diversity due to founder effect and changes in allele frequencies due to allele surfing at the expanding front(32).

Here we provide the first evidence that ALAN, alongside distance to broadleaf woodlands, affects genetic connectivity in *R. hipposideros* populations. While distance to woodlands was the main variable explaining genetic differentiation between colonies, ALAN also plays a role in limiting movement between roosts, potentially at a more localized scale, thereby influencing gene flow and functional connectivity. Our findings complement previous work on a light sensitive bat that avoids urban areas, the barbastelle bat, *Barbastella barbastellus(*33), augmenting our understanding of the wider influence of ALAN on landscape connectivity for nocturnal biodiversity.

In conclusion, using integrative approach through gene expression to landscape and adaptive ecology, our study provides the first evidence that ALAN induces stress in wild bat populations and influences longer-term genomic patterns at the population level reducing genetic connectivity with potential negative consequences for conservation. We demonstrate that ALAN can act as both an acute physiological stressor and contribute to evolutionary change. Consequently, as urban expansion and associated ALAN continue to intensify worldwide, the challenge for biodiversity conservation will be to not only mitigate their ecological impacts, but to also anticipate their evolutionary consequences. This will involve recognizing that while some species will be able to adapt to urbanization, others may shift their ranges, be subject to fitness related impacts and at threat of population decline without evidence-based interventions.

## Materials and Methods

### Experimental design and sampling

This study was approved by the University of Exeter Animal Welfare and Ethics Research Board. Work was carried out under licences from Natural England, Natural Resources Wales (license number 2023-66785-SCI-SCI) and the Home Office (PP0875087). We carried out lighting experiments at an *R. hipposideros* maternity roost in Somerset, England from 5^th^–9^th^ August 2024. We were not able to work on 8^th^ August due to bad weather. The first two nights served as the control (dark) treatment with no artificial light. The following two nights were the light treatment, during which a white LED floodlight (CORE Lighting, Stroud, UK) was deployed outside the roost entrance as bats were emerging for a maximum of 1 hour post dusk (Figure 1). Illumination at the point of emergence was monitored every 10 minutes throughout the survey period, using a T-10 illuminance meter (Konica Minolta Sensing Inc., Osaka, Japan). Mean illuminance was 0.9 lux on the control nights and 104 lux on the lit nights. We trapped bats as they emerged from the roost, sampling 10 bats per treatment group, spread over the four sampling nights (i.e. five bats per night) to minimize processing time.

Bats were captured with hand nets and harp traps following good practice guideline(34). To minimize biological variation in the gene expression analysis only non-lactating adult females were kept for sampling. Since *R. hipposideros* is a protected species in the UK, lab-based experiments and lethal sampling are not permitted. Blood was therefore chosen for examining changes in gene expression, as it is a proven non-lethal alternative for studying bat transcriptomes in wild populations(35).

Upon capture, bats were placed in individual cotton holding bags before processing. We recorded the age, reproductive status (non-breeding or post-lactating), forearm length and body mass of each bat. We then collected blood samples ensuring the total volume did not exceed 1% of each bat’s body mass (approximately 50 ul), which complies with the recommended limit for small mammals(36). We used sterilized insulin needles (32 gauge) to puncture the uropatagial vein and collected blood drops with a micropipette fitted with a sterile tip. Blood samples were immediately flash-frozen in liquid nitrogen on-site and transferred to -80^°^C freezers for storage prior to RNA extraction.

For whole genome sequencing (WGS), 19 sites were selected across England and Wales that represent varying levels of artificial light (Figure 1, Table S3). Bats were captured using the same techniques described above, either at dusk as the bats emerged, or during the daytime within the roost (by hand net only). Males, females and juveniles were included in the WGS sampling. A 3mm biopsy punch was collected from each wing of each bat and placed in absolute ethanol. We sampled 6-12 individuals per site, depending on roost size, to ensure we captured enough of the genetic variation present within a roost.

### RNA and DNA extraction and sequencing

Total RNA was extracted from blood using the NucleoSpin RNA Blood extraction kit (MACHEREY-NAGEL), per the manufacturer’s protocol. The yield and quality of RNA was evaluated using a Qubit Fluorometer 2.0 (Invitrogen) and a 4150 TapeStation system (Agilent). Libraries for each sample were built by the Exeter Sequencing Facility using the Illumina Stranded Total RNA Prep with Ribo-Zero. Sequencing was performed on a NovaSeq™ X Series using a paired-end approach with a read length of 150 bases. Total genomic DNA was extracted using the DNeasy Blood & Tissue Kit (Qiagen). The quantity and concentration of DNA was assessed by fluorometry using a Qubit 1X dsDNA High Sensitivity Assay Kit (Invitrogen). Libraries were prepared using the NEBnext UltraExpress FS DNA prep kit with NEB barcodes (8 bp UDIs). Sequencing was performed as described above, with a target of 10-15X sequencing depth per sample. Bioinformatics procedures are described in Supporting Information.

### Differential expression analysis

Differential expression was conducted using the Python package PyDESeq2 v0.5.0(37). Genes with less than 10 counts in total were omitted from downstream analysis. The DESeq2 model was run using treatment (light versus dark) as the design factor. Log fold change (LFC) shrinkage was implemented using the apeGLM prior. To minimize false-positives, genes were considered differentially expressed only if their adjusted *p*-values were less than 0.05 and their LFC values exceeded ±1. A principal components analysis (PCA) was performed on the differentially expressed genes (DEGs) using the normalized variance stabilizing transformation count matrix.

### Population structure and genomic diversity

A putatively neutral SNP dataset was created for analyses of population structure and landscape genetics. This dataset contained only SNPs that did not depart from Hardy-Weinberg equilibrium and were not identified as outliers by the R package OutFLANK v0.2(38). Population structure was assessed using non-model based principal component analysis (PCA) and model-based admixture inference using sparse non-negative matrix factorization (SNMF) from the R package LEA v3.12.2(39). For admixture analyses, SNMF was run independently ten times for each ancestral population (*K*). Cross-entropy statistics were extracted from the results and the optimal run for the *K* with the lowest cross-entropy was used for visualization of admixture (Figure S2). Individual admixture proportions were visualized as barplots, and the mean average proportion per cluster per location were visualized as admixture maps using mapmixture v1.2.0(40). Genetic differentiation between sites was assessed by computing pairwise *F*_ST_ values with gl.fst.pop() function from dartR v2.9.9(41) (Table S9). Two approaches were used to calculate genomic diversity. First, genome-wide heterozygosity (theta, excluding runs of homozygosity) and runs of homozygosity (ROH) were computed from the BAM alignments for each individual using ROHan v1.0.1(25). Second, observed heterozygosity (*H*_o_) was computed from the neutral SNP variants in R. To test for correlations between genetic diversity and landscape variables (described in next section), Spearman correlations were run where heterozygosity was modelled as a function of each environmental predictor variable.

### Landscape analysis

A landscape genetics approach(42) was used to assess the effect of ALAN and other landscape variables on gene flow and functional connectivity in *R. hipposideros*. We selected a set of landscape variables thought to affect genetic connectivity in these bats: extent of ALAN (downloaded from NASA Black Marble, https://blackmarble.gsfc.nasa.gov), Land Cover Map 2023 (UKCEH, www.ceh.ac.uk/data/ukceh-land-cover-maps), and Euclidean distance to woodland from roost (National Forest Inventory 2023, https://data-forestry.opendata.arcgis.com/). We converted landscape variables to resistance surfaces in R, assigning different resistance costs based on knowledge of the ecology of this species and its movement behavior. Resistance costs ranged from one, no resistance to movement, to 100, strong barrier to movement (Table S6), while the sea resistance was set to 200. We used Circuitscape v5.0.0(43) to calculate resistance distance matrices between *R. hipposideros* colonies based on the cumulative cost of movement due to landscape resistance.

We first used the R package ecodist(44) to test for the presence of isolation by distance (the tendency of geographically close roosts to be more genetically similar than more distant roosts due to dispersal limitations) by relating levels of genetic differentiation (*F*_ST_) to Euclidean distance between colonies. To account for isolation-by-distance, log *F*_ST_ (adding 1 to avoid negative values) was divided by log Euclidean distance. We then modelled pairwise log *F*_ST_ as a function of pairwise landscape resistances using the maximum likelihood population effects approach(45). Different combinations of landscape variables were modelled using the R package lme4 1.1-37(46) (Table S9), while the variance inflation factor (VIF<10) was calculated to remove correlated variables. Models were evaluated and compared using the AICc and BIC weighting criteria.

### Genotype-environment association (GEA)

We ran a genotype-environment association (GEA) analysis to identify genomic variants potentially associated with ALAN (NASA Black Marble) and urbanization (percent urban cover, UKCEH Land Cover Map 2023). All SNP variants were used in the GEA analysis. As GEA requires no missing data, genotypes were imputed with the impute() function from LEA using the most likely genotype (mode) and *K*=3 (number of ancestral populations), the *K* of which was informed by the population structure analysis. A partial redundancy analysis (pRDA) accounting for population structure was performed using the R package vegan v2.6-10 with the SNP genotypes as the dependent variable. The two environmental variables, ALAN and percent urban cover, were highly correlated (*r*^2^=0.93, *p*<0.001). Therefore, we ran a PCA on these two variables and used both principal component (PC) axes as predictor variables in the pRDA. GEA outliers, i.e. SNPs whose genotypes correlate significantly with the environmental predictors, were identified based on their extremeness along a distribution of Mahalanobis distances estimated between each SNP and the center of the RDA space using two axes(47). Bonferroni correction was used to account for multiple testing and SNPs with adjusted *p*-values <0.05 were considered as GEA outliers. The SNP positions on the chromosomes were then extracted and used to identify which GEA outliers are located directly in genes or within 1kb of a gene annotation.

### Biological pathway over-representation analysis

For the differentially expressed genes, biological pathway over-representation analysis was conducted using the Python package GSEApy v1.1.5(48). Gene set enrichment analysis (GSEA) is a computational method used to determine whether a set of DEGs are statistically over-represented in certain gene sets that are linked to biological functions, pathways or processes. Here, we used GSEApy as a wrapper to Enrichr(49), which requires a set of Entrez gene symbols as input to implement the GSEA. The following human gene set libraries were used in the analysis: GO Biological Process 2023 and MSigDB Hallmark 2020. GSEA was run using DEGs as the gene list and a background list of genes that passed filtering and thus were expressed in blood (*N*=16,519), and the libraries outlined above as the gene sets. The terms in each gene set were then filtered so that only terms with adjusted *p*-values less than 0.05 were considered enriched. The percentage of DEGs present in each gene set was also calculated to compare the proportion of DEGs identified across terms. For the genes identified as containing GEA outliers from the genotype-environment association analysis, the same Enrichr approach was also used, but for this analysis all the genes annotated in the *R. hipposideros* genome were used as the background list (*N*=18,890).

## Acknowledgments

This study was funded by the Natural Environment Research Council, grant number NE/W005778/1. We would like to thank the many bat groups, professionals and enthusiasts that helped us identify roost sites, and roost owners for allowing us access for trapping, particularly Ben McCarthy at the National Trust, Paula Tyler and Sam Smith. We thank Lewis Absalom, Pete Banfield, Nathan Biggs, Dan Flew, Olatz Gartzia, Jack Hooker, David Howes, Will Loach, William Rees, Rachel Reizin, Chiara Scaramella and Sam Smith for help with fieldwork. We also thank the Exeter Sequencing Facility, and in particular, Bryony Williams, Paul O’Neil, Audrey Farbos and Aaron Jeffries for support with the transcriptome and whole genome sequencing. We are grateful to Michael Hiller for facilitating access to the bat transcriptome.

## References

1. M. Morgan-Taylor, Regulating light pollution: More than just the night sky. Science 380, 1118–1120 (2023).

2. A. C. S. Owens, et al., Light pollution is a driver of insect declines. Biological Conservation 241, 108259 (2020).

3. C. C. M. Kyba, Y. Ö. Altıntaş, C. E. Walker, M. Newhouse, Citizen scientists report global rapid reductions in the visibility of stars from 2011 to 2022. Science 379, 265–268 (2023).

4. A. K. Jägerbrand, K. Spoelstra, Effects of anthropogenic light on species and ecosystems. Science 380, 1125–1130 (2023).

5. D. M. Dominoni, et al., Integrated molecular and behavioural data reveal deep circadian disruption in response to artificial light at night in male Great tits (Parus major). Sci Rep 12, 1553 (2022).

6. E. Knop, et al., Artificial light at night as a new threat to pollination. Nature 548, 206–209 (2017).

7. L. Eberhardt, H. B. Doria, B. Bulut, B. Feldmeyer, M. Pfenninger, Transcriptomics predicts artificial light at night’s (ALAN) negative fitness effects and altered gene expression patterns in the midge Chironomus riparius (Diptera:Chironomidae). Environmental Pollution 369, 125827 (2025).

8. D. M. Dominoni, J. Kjellberg Jensen, M. de Jong, M. E. Visser, K. Spoelstra, Artificial light at night, in interaction with spring temperature, modulates timing of reproduction in a passerine bird. Ecological Applications 30, e02062 (2020).

9. B. M. Van Doren, et al., High-intensity urban light installation dramatically alters nocturnal bird migration. Proceedings of the National Academy of Sciences 114, 11175–11180 (2017).

10. F.-S. Zhang, et al., Effects of artificial light at night on foraging behavior and vigilance in a nocturnal rodent. Science of The Total Environment 724, 138271 (2020).

11. L. A. Ramírez-Fráncel, et al., Bats and their vital ecosystem services: a global review. Integrative Zoology 17, 2–23 (2022).

12. E. L. Stone, S. Harris, G. Jones, Impacts of artificial lighting on bats: a review of challenges and solutions. Mammalian Biology 80, 213–219 (2015).

13. Y. Deng, et al., Behavioral and transcriptomic responses of wild least horseshoe bats to artificial light. iScience 28 (2025).

14. G. Jones, J. Rydell, Foraging strategy and predation risk as factors influencing emergence time in echolocating bats. Philos Trans R Soc Lond B Biol Sci 346, 445–455 (1994).

15. J. D. Hale, A. J. Fairbrass, T. J. Matthews, G. Davies, J. P. Sadler, The ecological impact of city lighting scenarios: exploring gap crossing thresholds for urban bats. Global Change Biology 21, 2467–2478 (2015).

16. E. C. Teeling, et al., Bat Biology, Genomes, and the Bat1K Project: To Generate Chromosome-Level Genomes for All Living Bat Species. Annual Review of Animal Biosciences 6, 23–46 (2018).

17. E. L. Stone, G. Jones, S. Harris, Street Lighting Disturbs Commuting Bats: Current Biology. Current Biology 19, 1123–1127 (2009).

18. J. Hildesheim, et al., Gadd45a Protects against UV Irradiation-induced Skin Tumors, and Promotes Apoptosis and Stress Signaling via MAPK and p5312. Cancer Res 62, 7305–7315 (2002).

19. J. Hu, C. M. McCall, T. Ohta, Y. Xiong, Targeted ubiquitination of CDT1 by the DDB1– CUL4A–ROC1 ligase in response to DNA damage. Nat Cell Biol 6, 1003–1009 (2004).

20. N. Aït-Ali, et al., Rod-Derived Cone Viability Factor Promotes Cone Survival by Stimulating Aerobic Glycolysis. Cell 161, 817–832 (2015).

21. I. Hernández-Muñoz, et al., Stable X chromosome inactivation involves the PRC1 Polycomb complex and requires histone MACROH2A1 and the CULLIN3/SPOP ubiquitin E3 ligase. Proceedings of the National Academy of Sciences 102, 7635–7640 (2005).

22. K. Rothbarth, et al., Induction of apoptosis by overexpression of the DNA-binding and DNA-PK-activating protein C1D. J Cell Sci 112, 2223–2232 (1999).

23. A. Takeuchi, et al., A Human Erythrocyte-derived Growth-promoting Factor with a Wide Target Cell Spectrum: Identification as Catalase1. Cancer Res 55, 1586–1589 (1995).

24. Y.-P. Wang, et al., Regulation of G6PD acetylation by SIRT2 and KAT9 modulates NADPH homeostasis and cell survival during oxidative stress. EMBO J 33, 1304–1320 (2014).

25. G. Renaud, K. Hanghøj, T. S. Korneliussen, E. Willerslev, L. Orlando, Joint Estimates of Heterozygosity and Runs of Homozygosity for Modern and Ancient Samples. Genetics 212, 587–614 (2019).

26. C. Vergata, et al., Transcriptome-wide deregulation of gene expression in zebrafish exposed to artificial light at night. Environmental Pollution 382, 126683 (2025).

27. E. V. Todd, M. A. Black, N. J. Gemmell, The power and promise of RNA-seq in ecology and evolution. Molecular Ecology 25, 1224–1241 (2016).

28. F. Thiebaut, A. S. Hemerly, P. C. G. Ferreira, A Role for Epigenetic Regulation in the Adaptation and Stress Responses of Non-model Plants. Front Plant Sci 10, 246 (2019).

29. A. M. Stankiewicz, A. H. Swiergiel, P. Lisowski, Epigenetics of stress adaptations in the brain. Brain Research Bulletin 98, 76–92 (2013).

30. L. Lehnen, et al., Genetic diversity in a long-lived mammal is explained by the past’s demographic shadow and current connectivity. Molecular Ecology 30, 5048–5063 (2021).

31. M. Veith, N. Beer, A. Kiefer, J. Johannesen, A. Seitz, The role of swarming sites for maintaining gene flow in the brown long-eared bat (Plecotus auritus). Heredity 93, 342–349 (2004).

32. L. Excoffier, N. Ray, Surfing during population expansions promotes genetic revolutions and structuration. Trends Ecol Evol 23, 347–351 (2008).

33. O. Razgour, et al., Applying genomic approaches to identify historic population declines in European forest bats. Journal of Applied Ecology 61, 160–172 (2024).

34. The Bat ConservationTrust, Bat Surveys for Professional Ecologists: Good Practice Guidelines 4th edition, 4th Ed. (The Bat Conservation Trust, 2023).

35. Z. Huang, et al., A nonlethal sampling method to obtain, generate and assemble whole blood transcriptomes from small, wild mammals. Molecular Ecology Resources 16, 150–162 (2016).

36. S. Parasuraman, R. Raveendran, R. Kesavan, Blood sample collection in small laboratory animals. J Pharmacol Pharmacother 1, 87–93 (2010).

37. B. Muzellec, M. Teleńczuk, V. Cabeli, M. Andreux, PyDESeq2: a python package for bulk RNA-seq differential expression analysis. Bioinformatics 39, btad547 (2023).

38. M. C. Whitlock, K. E. Lotterhos, Reliable Detection of Loci Responsible for Local Adaptation: Inference of a Null Model through Trimming the Distribution of F(ST). Am Nat 186 Suppl 1, S24–36 (2015).

39. C. Gain, O. François, LEA 3: Factor models in population genetics and ecological genomics with R. Mol Ecol Resour 21, 2738–2748 (2021).

40. T. L. Jenkins, mapmixture: An R package and web app for spatial visualisation of admixture and population structure. Mol Ecol Resour 24, e13943 (2024).

41. B. Gruber, P. J. Unmack, O. F. Berry, A. Georges, dartr: An r package to facilitate analysis of SNP data generated from reduced representation genome sequencing. Mol Ecol Resour 18, 691–699 (2018).

42. S. Manel, M. K. Schwartz, G. Luikart, P. Taberlet, Landscape genetics: combining landscape ecology and population genetics. Trends in Ecology & Evolution 18, 189–197 (2003).

43. B. H. McRae, B. G. Dickson, T. H. Keitt, V. B. Shah, Using Circuit Theory to Model Connectivity in Ecology, Evolution, and Conservation. Ecology 89, 2712–2724 (2008).

44. S. C. Goslee, D. L. Urban, The ecodist Package for Dissimilarity-based Analysis of Ecological Data. Journal of Statistical Software 22, 1–19 (2007).

45. M. J. Van Strien, D. Keller, R. Holderegger, A new analytical approach to landscape genetic modelling: least-cost transect analysis and linear mixed models. Molecular Ecology 21, 4010–4023 (2012).

46. D. Bates, M. Mächler, B. Bolker, S. Walker, Fitting Linear Mixed-Effects Models Using lme4. Journal of Statistical Software 67, 1–48 (2015).

47. T. Capblancq, B. R. Forester, Redundancy analysis: A Swiss Army Knife for landscape genomics. Methods in Ecology and Evolution 12, 2298–2309 (2021).

48. Z. Fang, X. Liu, G. Peltz, GSEApy: a comprehensive package for performing gene set enrichment analysis in Python. Bioinformatics 39, btac757 (2022).

49. Z. Xie, et al., Gene Set Knowledge Discovery with Enrichr. Curr Protoc 1, e90 (2021).

