## Supplementary material for "Artificial light at night induces stress and affects evolutionary change in bats": Table S1; Figure S1

**Supporting Information for**
### Artificial light at night induces stress and affects evolutionary change in bats

Tom L. Jenkins, Thomas Foxley, Yury Malovichko, Emma L. Stone, Orly Razgour*

**This PDF file includes:**

Supporting methods
Figures S1 to S2
Tables S1 to S9
SI References

#### Supporting Methods

**Extended description of bioinformatics procedures**

FASTQ files were trimmed using fastp v23.4(1) with the following parameters: (i) polyG and polyX trimming enabled, (ii) a Phred quality score of 30 or more in at least 60% of bases in a read, and (iii) a minimum length of 100bp. For RNA sequencing, Kallisto v0.51.1(2) was used to quantify transcripts in each sample using the trimmed reads and the R. hipposideros transcriptome coding sequences (GCA_964194185.1; HLrhiHip1A). A custom Python script (see Data Archiving Statement) was written to amalgamate raw counts for each sample into a single count matrix where each column represented a sample and each row represented a gene. For WGS, reads were aligned to the R. hipposideros genome (GCA_964194185.1) using Bowtie2 v2.5.3(3) and BAM files were processed (adding read groups, marking duplicates and sorting by coordinate) using Samtools v1.19(4). Variants were called in parallel using Freebayes v1.3.8(5) with ploidy set to 2, minimum base quality set to 20, minimum mapping quality set to 20 and a maximum coverage set to 1000X. Single nucleotide polymorphisms (SNPs) were filtered in the following order: (1) to reduce the dataset, a 20% maximum missing genotypes and a minor allele count threshold >= was enforced; (2) removal of two individuals with extremely high missing data (>50% missing); and (3) a final filter of 10% maximum missing genotypes, genotype qualities >= 15, a minor allele count >= 3, and thinning of SNPs within 1kb to reduce linkage disequilibrium.

#### Figures

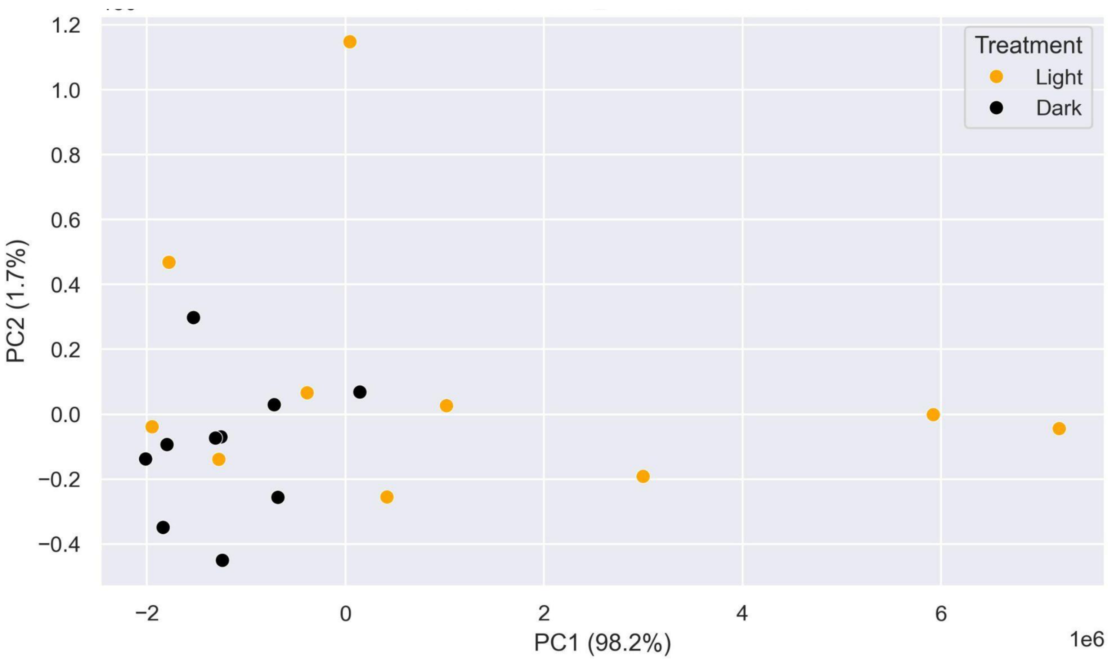

**Fig. S1.** Principal components analysis (PCA) on the normalized count matrix for 252 differentially expressed genes (DEGs). Each point represents an individual sample, and the colors denote whether a bat was exposed to artificial light (orange) or was not exposed and are considered dark controls (black).

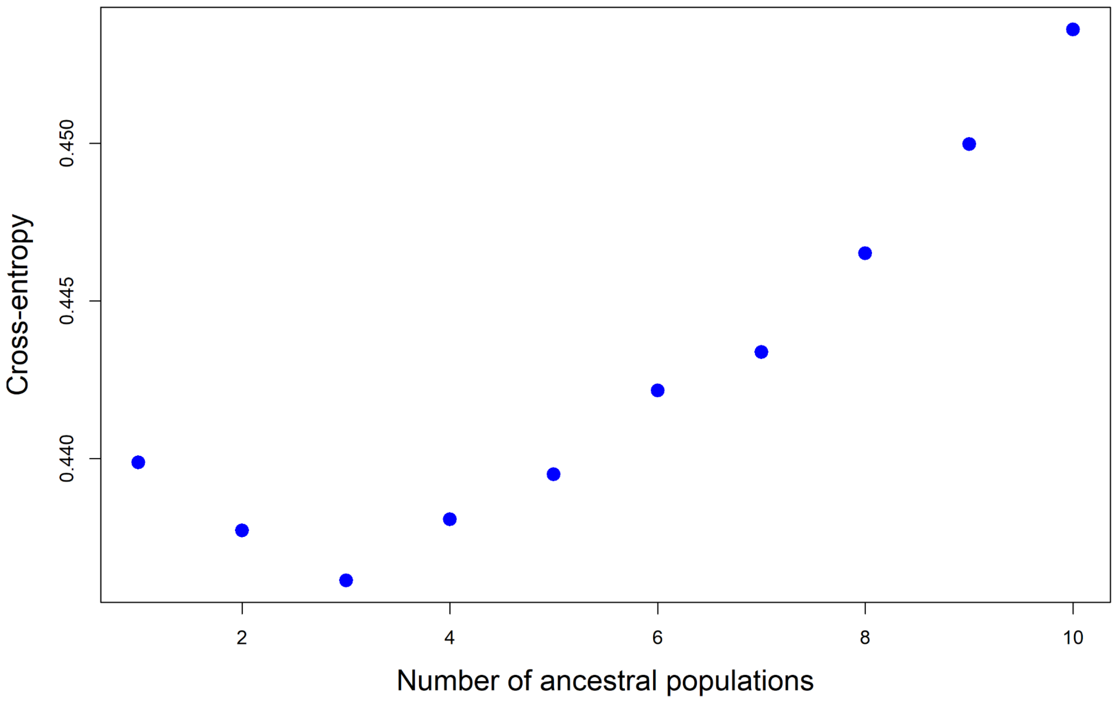

**Fig. S2.** Sparse non-negative matrix factorization (SNMF) cross-entropy results. Lower cross-entropy values indicate that a particular K (ancestral populations) has higher statistical support. An ‘elbow point’ is clearly observed at K=3, so K=3 was used in the interpretation and visualization of admixture (ancestry).

#### Tables

**Table S1.** Differentially expressed genes in Rhinolophus hipposideros exposed to artificial lights.

| Differentially Expressed Gene (DEG) | log2 Fold Change (LFC) | Adjust P value | Reported Roles and Functions | References |
|---|---|---|---|---|
| IFI27 | 2.09 | 0.007 | Apoptosis. | https://www.uniprot.org/uniprotkb/P40305 |
| HBB | 1.77 | 0.007 | Oxygen transport. | https://www.uniprot.org/uniprotkb/P68871 |
| VPS28 | 1.44 | 0.007 | Regulator of vesicular trafficking process. | https://www.uniprot.org/uniprotkb/Q9UK41 |
| ST3GAL4 | 1.32 | 0.007 | Haemostasis (stopping the flow of blood). | https://www.uniprot.org/uniprotkb/Q11206/ |
| COPS5 | 1.18 | 0.007 | Regulator of ubiquitin conjugation pathway. | https://www.uniprot.org/uniprotkb/Q92905 |
| CMAS | 1.45 | 0.008 | Amino-sugar metabolism. | https://www.uniprot.org/uniprotkb/Q8NFW8 |
| TSPO2 | 1.64 | 0.012 | Role in transport processes at the plasma membrane of erthrocytes. | https://www.uniprot.org/uniprotkb/Q5TGU0 |
| MED30 | 1.22 | 0.012 | Transcription regulation. | https://www.uniprot.org/uniprotkb/Q96HR3 |
| TMEM256 | 1.46 | 0.012 | Transmembrane protein. | |
| MPLKIP | 1.17 | 0.012 | Maintenance of cell cycle integrity by regulating mitosis or cytokinesis. | https://www.uniprot.org/uniprotkb/Q8TAP9 |
| N4BP2L1 | 1.64 | 0.012 | | |
| UHRF1 | 1.50 | 0.012 | Maintenance of DNA methylation, chromatin modification and DNA repair. | https://www.uniprot.org/uniprotkb/Q96T88 |
| ESCO2 | 1.38 | 0.012 | DNA replication. | https://www.uniprot.org/uniprotkb/Q56NI9 |
| HADHB | 1.37 | 0.012 | Metabolism. | https://www.uniprot.org/uniprotkb/P55084 |
| C4BPA | 1.53 | 0.015 | | |
| MGLL | 1.46 | 0.015 | Metabolism. | https://www.uniprot.org/uniprotkb/Q99685 |
| MACROD1 | 1.44 | 0.015 | Involved in DNA damage response | https://www.ncbi.nlm.nih.gov/gene/28992 |
| CARHSP1 | 1.38 | 0.015 | | |
| ATP5IF1 | 1.38 | 0.015 | Regulates heme synthesis by modulating the mitochondrial pH and redox potential. | https://www.uniprot.org/uniprotkb/Q9UII2 |
| DCK | 1.34 | 0.015 | Phosphorylation. | https://www.ncbi.nlm.nih.gov/gene/1633 |

| | | | | |
|---|---|---|---|---|
| NVL | 1.28 | 0.015 | Telomerase activity. | https://www.uniprot.org/uniprotkb/O15381/entry |
| GHITM | 1.27 | 0.015 | Maintenance of mitochondrial morphology. | https://www.uniprot.org/uniprotkb/Q9H3K2/entry |
| TRIM23 | 1.18 | 0.015 | Autophagy activation. | https://www.uniprot.org/uniprotkb/P36406/entry |
| KPTN | 1.16 | 0.015 | | |
| SMIM14 | 1.13 | 0.015 | | |
| NUP133 | 1.02 | 0.015 | RNA transport. | https://www.uniprot.org/uniprotkb/Q8WUM0/entry |
| PSMD14 | 1.01 | 0.015 | Maintenance of protein homeostasis. | https://www.uniprot.org/uniprotkb/O00487/entry |
| FAM228A | 1.83 | 0.016 | | |
| CD68 | 1.46 | 0.016 | | |
| PRSS27 | 1.44 | 0.016 | | |
| SNCA | 1.36 | 0.016 | Synaptic activity. | https://www.uniprot.org/uniprotkb/P37840/entry |
| UBE2B | 1.36 | 0.016 | DNA repair. | https://www.uniprot.org/uniprotkb/P63146/entry |
| CXorf38 | 1.31 | 0.016 | | |
| TTC7B | 1.31 | 0.016 | | |
| MACROH2A1 | 1.28 | 0.016 | Transcription regulation, DNA repair, DNA replication and chromosomal stability. | https://www.uniprot.org/uniprotkb/O75367/entry |
| WDR45 | 1.28 | 0.016 | Intracellular degradation process. | https://www.uniprot.org/uniprotkb/Q9Y484/entry |
| MAP1LC3B | 1.27 | 0.016 | Cellular stress. Mitochondrial metabolism. | https://www.uniprot.org/uniprotkb/Q9GZQ8/entry |
| CREG1 | 1.26 | 0.016 | Transcriptional control of cell growth and differentiation. | https://www.uniprot.org/uniprotkb/O75629/entry |
| RNF144A | 1.23 | 0.016 | DNA damage response. | https://www.uniprot.org/uniprotkb/P50876/entry |
| FAXDC2 | 1.22 | 0.016 | Promotes megakaryocyte differentiation (blood clotting). | https://www.uniprot.org/uniprotkb/Q96IV6/entry |
| TMC8 | 1.20 | 0.016 | Regulation of cellular processes. | https://www.uniprot.org/uniprotkb/Q8IU68/entry |
| RBX1 | 1.17 | 0.016 | cell cycle progression, signal transduction, transcription and transcription-coupled nucleotide excision repair | https://www.uniprot.org/uniprotkb/P62877/entry |
| C1D | 1.15 | 0.016 | Can induce apoptosis. | https://www.uniprot.org/uniprotkb/Q13901/entry |
| ATP5PB | 1.10 | 0.016 | Mitochondrial ATP synthase | https://www.uniprot.org/uniprotkb/P24539/entry |
| CHAC2 | 1.09 | 0.016 | | |
| ISG15 | 1.05 | 0.016 | | |

| | | | | |
|---|---|---|---|---|
| CLEC20A | 1.40 | 0.016 | | |
| PSMF1 | 1.34 | 0.016 | Control of proteasome function. | https://www.uniprot.org/uniprotkb/Q92530/entry |
| DDB1 | 1.33 | 0.016 | Part of the UV-DDB complex, plays a crucial role in recognizing and initiating repair of UV-induced DNA damage. | https://www.uniprot.org/uniprotkb/Q16531 |
| RAB27B | 1.33 | 0.016 | Homeostasis. | https://www.uniprot.org/uniprotkb/O00194/entry |
| HBD | 1.30 | 0.016 | Oxygen transport. | https://www.uniprot.org/uniprotkb/P02042/entry |
| FGD6 | 1.30 | 0.016 | | |
| PDK2 | 1.29 | 0.016 | Metabolism and homeostasis. Plays an important role in maintaining normal blood glucose levels and in metabolic adaptation to nutrient availability. Plays a role in the regulation of cell proliferation and in resistance to apoptosis under oxidative stress. Plays a role in p53/TP53-mediated apoptosis. | https://www.uniprot.org/uniprotkb/Q15119/entry |
| FBN1 | 1.28 | 0.016 | Tissue homeostasis. | https://www.uniprot.org/uniprotkb/P35555/entry |
| MPP1 | 1.27 | 0.016 | Essential regulator of neutrophil polarity. | https://www.uniprot.org/uniprotkb/Q00013/entry |
| H2BC11 | 1.26 | 0.016 | Histones thereby play a central role in transcription regulation, DNA repair, DNA replication and chromosomal stability. | https://www.uniprot.org/uniprotkb/P06899/entry |
| BLVRB | 1.26 | 0.016 | | |
| RNF38 | 1.26 | 0.016 | | |
| HSPH1 | 1.26 | 0.016 | Heat shock protein. | https://www.uniprot.org/uniprotkb/Q92598/entry |
| TCP11L2 | 1.24 | 0.016 | | |
| TAX1BP1 | 1.22 | 0.016 | | |
| GABARAPL2 | 1.21 | 0.016 | Role in mitophagy. | https://www.uniprot.org/uniprotkb/P60520/entry |
| GSTP1 | 1.21 | 0.016 | | |
| ELL2 | 1.21 | 0.016 | | |
| USP15 | 1.20 | 0.016 | DNA repair. | https://www.uniprot.org/uniprotkb/Q9Y4E8/entry |
| DEPDC1 | 1.20 | 0.016 | | |
| H3-3A | 1.20 | 0.016 | | |
| ATP13A2 | 1.20 | 0.016 | Mitochondrial ATP synthase | |
| LSM12 | 1.20 | 0.016 | | |

| | | | | |
|---|---|---|---|---|
| SIRT2 | 1.20 | 0.016 | Acts as a key regulator in the pentose phosphate pathway (PPP) by deacetylating and activating the glucose-6-phosphate G6PD enzyme, and therefore, stimulates the production of cytosolic NADPH to counteract oxidative damage. | https://www.uniprot.org/uniprotkb/Q8IXJ6/entry |
| PARK7 | 1.19 | 0.016 | Multifunctional protein with controversial molecular function which plays an important role in cell protection against oxidative stress and cell death acting as oxidative stress sensor and redox-sensitive chaperone and protease. | https://www.uniprot.org/uniprotkb/Q99497/entry |
| MPG | 1.19 | 0.016 | | |
| SEC62 | 1.19 | 0.016 | | |
| TRIM10 | 1.19 | 0.016 | | |
| MAP3K20 | 1.18 | 0.016 | | |
| ATP5PF | 1.18 | 0.016 | Mitochondrial ATP synthase | |
| MAD2L1 | 1.18 | 0.016 | | |
| JPT1 | 1.17 | 0.016 | | |
| HK1 | 1.17 | 0.016 | | |
| CHMP5 | 1.17 | 0.016 | | |
| SMC2 | 1.16 | 0.016 | | |
| WRAP53 | 1.15 | 0.016 | | |
| UBE2L6 | 1.15 | 0.016 | | |
| SELENOK | 1.13 | 0.016 | | |
| VDAC3 | 1.10 | 0.016 | | |
| ACO2 | 1.10 | 0.016 | | |
| RNPS1 | 1.10 | 0.016 | | |
| RBBP4 | 1.10 | 0.016 | | |
| ZFAND3 | 1.05 | 0.016 | | |
| RRAGB | 1.05 | 0.016 | | |
| MOSPD1 | 1.05 | 0.016 | | |
| SOD1 | 1.02 | 0.016 | Superoxide dismutase, crucial for detoxifying superoxide radicals. Destroys radicals which are normally produced within the cells and which are toxic to biological systems. | https://www.uniprot.org/uniprotkb/P00441/entry |
| FUNDC2 | 1.01 | 0.016 | | |
| HSD17B10 | 1.00 | 0.016 | | |
| LAMTOR5 | 1.00 | 0.016 | | |
| H2AC11 | 1.19 | 0.016 | | |
| SLC1A5 | 1.17 | 0.016 | | |
| RIOK3 | 1.18 | 0.016 | | |
| PTPRN | 1.15 | 0.016 | | |
| ANP32E | 1.16 | 0.017 | | |

| | | |
|---|---|---|
| DNAJC8 | 1.10 | 0.017 |
| TSPAN33 | 1.35 | 0.017 |
| RABAC1 | 1.09 | 0.017 |
| TMEM86B | 1.37 | 0.017 |
| RHAG | 1.20 | 0.017 |
| EIF2AK1 | 1.19 | 0.017 |
| CDKN2D | 1.12 | 0.017 |
| COPZ1 | 1.11 | 0.017 |
| GPR19 | 1.11 | 0.017 |
| C6orf226 | 1.11 | 0.017 |
| ISCA1 | 1.10 | 0.017 |
| AP2B1 | 1.06 | 0.017 |
| FRRS1 | 1.17 | 0.017 |
| PSMC3 | 1.17 | 0.017 |
| COX7B | 1.15 | 0.017 |
| UROS | 1.26 | 0.017 |
| TFDP2 | 1.11 | 0.017 |
| REEP1 | 1.03 | 0.017 |
| GPD2 | 1.40 | 0.017 |
| ATOX1 | 1.20 | 0.017 |
| ERMAP | 1.19 | 0.017 |
| SLC25A11 | 1.16 | 0.017 |
| CCNI | 1.14 | 0.017 |
| CTSD | 1.11 | 0.017 |
| RPL35 | 1.11 | 0.017 |
| ST13 | 1.09 | 0.017 |
| UBA52 | 1.08 | 0.017 |
| BRCC3 | 1.05 | 0.017 |
| RPS14 | 1.04 | 0.017 |
| CALCOCO2 | 1.04 | 0.017 |
| COX6B1 | 1.03 | 0.017 |
| CLIC2 | 1.01 | 0.017 |
| SLC4A1 | 1.27 | 0.017 |
| KCND1 | 1.27 | 0.017 |
| CCDC34 | 1.27 | 0.017 |
| RNF123 | 1.25 | 0.017 |
| CALR | 1.17 | 0.017 |
| SYAP1 | 1.12 | 0.017 |
| IFT22 | 1.06 | 0.017 |
| COPG1 | 1.05 | 0.017 |
| NDUFA4 | 1.00 | 0.017 |
| SLC22A4 | 1.26 | 0.017 |
| MBNL3 | 1.14 | 0.017 |
| PCBP1 | 1.05 | 0.017 |
| GNLY | 1.95 | 0.017 |

| | | | | |
|---|---|---|---|---|
| DCN | 1.54 | 0.017 | | |
| RSAD2 | 1.31 | 0.017 | | |
| AP2A1 | 1.24 | 0.017 | | |
| DENND4A | 1.18 | 0.017 | | |
| TMEM98 | 1.18 | 0.017 | | |
| AZIN1 | 1.15 | 0.017 | | |
| SKP1 | 1.14 | 0.017 | | |
| RAD23A | 1.14 | 0.017 | | |
| NDUFV1 | 1.13 | 0.017 | | |
| GABARAP | 1.12 | 0.017 | | |
| GYPC | 1.07 | 0.017 | | |
| GLO1 | 1.06 | 0.017 | | |
| TBL1XR1 | 1.06 | 0.017 | | |
| GSTA4 | 1.04 | 0.017 | | |
| RPS9 | 1.04 | 0.017 | | |
| PCK2 | 1.04 | 0.017 | | |
| RB1 | 1.04 | 0.017 | | |
| UQCRH | 1.03 | 0.017 | | |
| PTP4A2 | 1.01 | 0.017 | | |
| CAT | 1.29 | 0.017 | Catalyzes the degradation of hydrogen peroxide (H2O2) generated by peroxisomal oxidases to water and oxygen, thereby protecting cells from the toxic effects of hydrogen peroxide. | https://www.uniprot.org/uniprotkb/P04040/entry |
| CLEC3B | 1.41 | 0.018 | | |
| RANBP10 | 1.17 | 0.018 | | |
| GATA1 | 1.16 | 0.018 | | |
| SLC46A3 | 1.10 | 0.018 | | |
| FBXO9 | 1.09 | 0.018 | | |
| ADD2 | 1.27 | 0.018 | | |
| PRXL2A | 1.13 | 0.018 | | |
| RNASEH2C | 1.13 | 0.018 | | |
| KEL | 1.13 | 0.018 | | |
| DCUN1D1 | 1.06 | 0.018 | | |
| PSMD4 | 1.05 | 0.018 | | |
| MPC2 | 1.05 | 0.018 | | |
| HMOX2 | 1.04 | 0.018 | | |
| SLC43A1 | 1.04 | 0.018 | | |
| ZMAT5 | 1.04 | 0.018 | | |
| COX5B | 1.03 | 0.018 | | |
| ACSS1 | 1.02 | 0.018 | | |
| AP2S1 | 1.01 | 0.018 | | |
| ST3GAL5 | 1.06 | 0.018 | | |
| NFIX | 1.08 | 0.018 | | |

| | | | | |
|---|---|---|---|---|
| RAB31 | 1.14 | 0.018 | | |
| BSG | 1.05 | 0.018 | Retinal maturation and development. Acts as a retinal cell surface receptor for NXNL1 and plays an important role in NXNL1-mediated survival of retinal cone photoreceptors. | https://www.uniprot.org/uniprotkb/P35613/entry |
| ZFAND4 | 1.10 | 0.018 | | |
| GLRB | 1.21 | 0.018 | | |
| DMTN | 1.15 | 0.018 | | |
| WBP2 | 1.10 | 0.018 | | |
| PAQR5 | 1.05 | 0.018 | | |
| RGS22 | 1.04 | 0.018 | | |
| TMBIM6 | 1.03 | 0.018 | | |
| TPT1 | 1.02 | 0.018 | | |
| NAA80 | 1.09 | 0.018 | | |
| RPLP0 | 1.11 | 0.019 | | |
| GOLPH3L | 1.08 | 0.019 | | |
| PABPC1 | 1.07 | 0.019 | | |
| AP1B1 | 1.03 | 0.019 | | |
| PLEKHA6 | 1.15 | 0.019 | | |
| STRADB | 1.13 | 0.019 | | |
| HYAL3 | 1.12 | 0.019 | | |
| ASIC4 | 1.17 | 0.019 | | |
| CA2 | 1.08 | 0.019 | | |
| RPLP1 | 1.06 | 0.019 | | |
| SRXN1 | 1.04 | 0.019 | Contributes to oxidative stress resistance by reducing cysteine-sulfinic acid formed under exposure to oxidants in the peroxiredoxins PRDX1, PRDX2, PRDX3 and PRDX4. | https://www.uniprot.org/uniprotkb/Q9BYN0/entry |
| TRAK2 | 1.02 | 0.019 | | |
| TIMM17B | 1.01 | 0.019 | | |
| GADD45A | 1.01 | 0.019 | Gadd45a is a critical factor protecting the epidermis against UV radiation-induced tumorigenesis by promoting damaged keratinocytes to undergo apoptosis and/or cell cycle arrest, two crucial events that prevent the expansion of mutant or deregulated cells. | https://pubmed.ncbi.nlm.nih.gov/12499274/#:~:text=Herein%20we%20demonstrate%20that%20Gadd45a,skin%20against%20UV%2Dinduced%20tumors. |
| PLEK2 | 1.01 | 0.019 | | |
| DOP1B | 1.14 | 0.019 | | |
| MAGIX | 1.06 | 0.019 | | |

| | | | | |
|---|---|---|---|---|
| PRDX2 | 1.04 | 0.019 | Thiol-specific peroxidase that catalyzes the reduction of hydrogen peroxide and organic hydroperoxides to water and alcohols, respectively. Plays a role in cell protection against oxidative stress by detoxifying peroxides and as sensor of hydrogen peroxide-mediated signalling events. | https://www.uniprot.org/uniprotkb/P32119/entry |
| TRPM6 | 1.02 | 0.019 | | |
| MTFR1 | 1.08 | 0.020 | | |
| SPTB | 1.16 | 0.020 | | |
| CANX | 1.01 | 0.020 | | |
| CALCOCO1 | 1.08 | 0.020 | | |
| DYNLL1 | 1.03 | 0.020 | | |
| MOB1B | 1.02 | 0.020 | | |
| SLC24A1 | 1.37 | 0.020 | | |
| PHOSPHO1 | 1.18 | 0.020 | | |
| YBX3 | 1.04 | 0.020 | | |
| XK | 1.03 | 0.020 | | |
| PTGS2 | -1.54 | 0.020 | | |
| OR2W3 | 1.11 | 0.020 | | |
| MXI1 | 1.03 | 0.020 | | |
| ACSL1 | 1.01 | 0.020 | | |
| SNX9 | 1.00 | 0.020 | | |
| SMOX | 1.14 | 0.021 | | |
| ISCA2 | 1.01 | 0.021 | | |
| NCEH1 | 1.13 | 0.021 | | |
| GCH1 | 1.09 | 0.021 | | |
| RPS2 | 1.05 | 0.021 | | |
| TFRC | 1.04 | 0.021 | | |
| GLUL | 1.04 | 0.021 | | |
| SCARB1 | 1.04 | 0.021 | | |
| GPX1 | 1.02 | 0.021 | Catalyzes the reduction of hydroperoxides. | https://www.uniprot.org/uniprotkb/P07203 |
| FKBP8 | 1.01 | 0.021 | | |
| SLC25A37 | 1.00 | 0.021 | | |
| SLC6A9 | 1.12 | 0.021 | | |
| MSMB | 1.25 | 0.021 | | |
| FAM214B | 1.07 | 0.021 | | |
| FBXO48 | 1.13 | 0.022 | | |
| PAQR9 | 1.13 | 0.022 | | |
| PLCB1 | 1.06 | 0.022 | | |
| HEMGN | 1.38 | 0.023 | | |
| UBE2O | 1.13 | 0.023 | | |
| UGGT2 | 1.06 | 0.024 | | |

| | | |
|---|---|---|
| RNF19B | 1.01 | 0.025 |
| PTPN14 | 1.03 | 0.026 |

**Table S2.** Gene set enrichment analysis results for differentially expressed genes in Rhinolophus hipposideros exposed to artificial lights (GO = GO Biological Process 2023, MSigDB = MSigDB Hallmark 2020).

| Gene Set | Term | Adjusted P value | Odds Ratio | Combined Score | Differentially Expressed Genes in Gene Set | Percent DEGs in Gene Set (%) |
|---|---|---|---|---|---|---|
| GO | Carbon Dioxide Transport (GO:0015670) | 0.002 | 110.5 | 1481.4 | CA2;HBB;RHAG;HBD | 36.4 |
| MSigDB | heme Metabolism | 0.000 | 19.0 | 1477.3 | SLC22A4;ISCA1;USP15;TFRC;GYPC;DCUN1D1;HBB;MOSPD1;SLC4A1;HBD;GATA1;TRAK2;ADD2;PRDX2;CA2;BSG;MXI1;CLIC2;ERMAP;FBXO9;SNCA;MPP1;DMTN;RIOK3;SMOX;EIF2AK1;UROS;RAD23A;ELL2;SPTB;RNF123;SLC25A37;TSPO2;KEL;XK;SLC6A9;TFDP2;CAT;RANBP10;BLVRB;RHAG;TRIM10 | 21.0 |
| GO | Gas Transport (GO:0015669) | 0.006 | 44.2 | 500.2 | CA2;HBB;RHAG;HBD | 26.7 |
| GO | Hydrogen Peroxide Catabolic Process (GO:0042744) | 0.002 | 34.7 | 453.1 | PRDX2;GPX1;CAT;HBB;HBD | 25.0 |
| GO | Cellular Response To Nitrogen Levels (GO:0043562) | 0.024 | 41.3 | 353.1 | GABARAPL2;MAP1LC3B;GABARAP | 27.3 |
| GO | Cellular Response To Nitrogen Starvation (GO:0006995) | 0.024 | 41.3 | 353.1 | GABARAPL2;MAP1LC3B;GABARAP | 27.3 |
| GO | Mitochondrial Electron Transport, Cytochrome C To Oxygen (GO:0006123) | 0.016 | 24.6 | 236.6 | COX7B;NDUFA4;COX5B;COX6B1 | 25.0 |
| MSigDB | Reactive Oxygen Species Pathway | 0.000 | 11.2 | 146.8 | CDKN2D;PRDX2;SRXN1;CAT;HMOX2;ATOX1;SOD1;LAMTOR5 | 16.3 |
| GO | One-Carbon Compound Transport (GO:0019755) | 0.021 | 12.6 | 115.1 | CA2;HBB;SLC4A1;RHAG;HBD | 14.3 |
| GO | Amino Acid Import Across Plasma | 0.033 | 13.8 | 108.0 | SLC22A4;TSPO2;SLC6A9;SLC1A5 | 14.8 |

| | | | | | | |
|---|---|---|---|---|---|---|
| | Membrane (GO:0089718) | | | | | |
| GO | Mitochondrion Disassembly (GO:0061726) | 0.032 | 9.5 | 76.4 | FUNDC2;GABARAPL2;MAP1LC3B;WDR45;GABARAP | 13.5 |
| GO | Regulation Of Protein Localization To Nucleus (GO:1900180) | 0.023 | 8.5 | 75.4 | TFRC;NVL;ATP13A2;PARK7;GLUL;LAMTOR5 | 11.5 |
| GO | Substantia Nigra Development (GO:0021762) | 0.033 | 9.2 | 72.6 | ATP5PF;ATP5PB;DYNLL1;SIRT2;COX6B1 | 11.9 |
| GO | Negative Regulation Of Protein Catabolic Process (GO:0042177) | 0.024 | 8.1 | 69.7 | GABARAPL2;PSMF1;SIRT2;AZIN1;SNCA;MAD2L1 | 11.1 |
| GO | Energy Derivation By Oxidation Of Organic Compounds (GO:0015980) | 0.034 | 8.9 | 69.0 | COX7B;NDUFA4;COX5B;UQCRH;COX6B1 | 11.6 |
| GO | Erythrocyte Differentiation (GO:0030218) | 0.024 | 7.9 | 67.1 | ATP5IF1;RPS14;TSPO2;DMTN;SLC1A5;GATA1 | 12.2 |
| MSigDB | Fatty Acid Metabolism | 0.000 | 5.5 | 65.5 | HADHB;HSPH1;ACSL1;CA2;GPD2;UROS;UBE2L6;ACO2;ACSS1;GLUL;HSD17B10;MGLL | 7.6 |
| GO | Response To Reactive Oxygen Species (GO:0000302) | 0.021 | 7.1 | 64.0 | GPX1;PTPRN;GSTP1;CAT;HBB;PDK2;SOD1 | 10.6 |
| GO | Regulation Of Proteasomal Protein Catabolic Process (GO:0061136) | 0.029 | 7.6 | 62.3 | GABARAPL2;GPX1;PSMD14;PSMC3;RAD23A;PSMF1 | 11.3 |
| GO | Cellular Response To Chemical Stress (GO:0062197) | 0.017 | 6.4 | 60.3 | PRDX2;GPX1;SRXN1;ATP13A2;PARK7;SIRT2;RBX1;SNCA | 9.0 |
| GO | Regulation Of Protein Catabolic Process (GO:0042176) | 0.010 | 5.6 | 58.6 | DDB1;GPX1;PSMD14;C4BPA;ATP13A2;VPS28;AZIN1;SIRT2;RBX1;MAD2L1 | 8.2 |

| | | | | | | |
|---|---|---|---|---|---|---|
| MSig DB | Oxidative Phosphorylation | 0.000 | 4.4 | 49.7 | ATP5PF;ISCA1;COX7B;ATP5PB;NDUFA4;COX5B;HSD17B10;UQCRH;COX6B1;HADHB;VDAC3;ACO2;NDUFV1;SLC25A11 | 7.0 |
| GO | Aerobic Electron Transport Chain (GO:0019646) | 0.049 | 6.1 | 43.3 | COX7B;NDUFA4;COX5B;NDUFV1;UQCRH;COX6B1 | 8.8 |
| GO | Macroautophagy (GO:0016236) | 0.032 | 5.1 | 40.5 | GABARAPL2;MAP1LC3B;CALCOCO2;ATP13A2;CALR;VPS28;GABARAP;CHMP5 | 7.7 |
| GO | Cellular Respiration (GO:0045333) | 0.045 | 5.2 | 37.8 | COX7B;NDUFA4;MTFR1;COX5B;NDUFV1;UQCRH;COX6B1 | 8.2 |
| GO | Ubiquitin-Dependent Protein Catabolic Process (GO:0006511) | 0.006 | 3.4 | 37.6 | PSMD14;UBE2B;UHRF1;RNF19B;UBE2L6;RAD23A;SIRT2;RBX1;RNF144A;DDB1;IFI27;PSMC3;PSMD4;TBL1XR1;PSMF1;VPS28;CHMP5;SKP1;FBXO9 | 5.2 |
| GO | Receptor-Mediated Endocytosis (GO:0006898) | 0.045 | 4.5 | 32.7 | RAB31;TFRC;CANX;AP2S1;SNX9;AP2A1;AP2B1;SNCA | 6.6 |
| GO | Cellular Response To Oxidative Stress (GO:0034599) | 0.045 | 4.5 | 32.7 | PRDX2;GPX1;SRXN1;ATP13A2;PARK7;PDK2;SIRT2;SNCA | 6.8 |
| GO | Negative Regulation Of Apoptotic Process (GO:0043066) | 0.011 | 3.1 | 32.0 | CDKN2D;RB1;GPX1;TFRC;GSTP1;GLO1;PARK7;GATA1;SOD1;PRDX2;COPS5;CAT;TAX1BP1;FKBP8;TMBIM6;TPT1;SNCA;MAD2L1;LAMTOR5 | 3.9 |
| GO | Proteasomal Protein Catabolic Process (GO:0010498) | 0.032 | 3.5 | 27.8 | DDB1;PSMD4;IFI27;PSMC3;PSMD14;UBE2B;TBL1XR1;RAD23A;SIRT2;FBXO9;SKP1;RBX1 | 5.5 |
| GO | Modification-Dependent Protein Catabolic Process (GO:0019941) | 0.037 | 3.6 | 27.3 | DDB1;RNF144A;PSMD14;UBE2B;UHRF1;RNF19B;UBE2L6;ISG15;PSMF1;UBA52;RBX1 | 5.7 |
| MSig DB | Adipogenesis | 0.005 | 3.5 | 26.1 | SCARB1;COX7B;GADD45A;RIOK3;GPD2;CAT;GHITM;SLC1A5;ACO2;MGLL;SOD1 | 5.5 |
| MSig DB | Myc Targets V1 | 0.006 | 3.3 | 23.5 | COPS5;PSMD14;PCBP1;GLO1;RPLP0;CANX;VDAC3;RNPS1;PABPC1;RPS2;MAD2L1 | 5.5 |

| | | | | | | |
|---|---|---|---|---|---|---|
| MSig DB | UV Response Up | 0.020 | 3.4 | 18.7 | TFRC;GCH1;PSMC3;CA2;CREG1;BSG;AP2S1;TMBIM6 | 5.1 |
| MSig DB | Xenobiotic Metabolism | 0.019 | 3.2 | 18.4 | SLC46A3;GCH1;CA2;SMOX;CAT;BLVRB;TMBIM6;SLC1A5;ACO2 | 4.5 |
| GO | Regulation Of Apoptotic Process (GO:0042981) | 0.045 | 2.3 | 16.7 | TFRC;GADD45A;GSTP1;GLO1;PARK7;GATA1;SOD1;PRDX2;COPS5;CAT;TAX1BP1;MAP3K20;ANP32E;FKBP8;ATP13A2;RNPS1;TMBIM6;CALR;CTSD;TPT1;SNCA;MAD2L1 | 3.1 |

**Table S3.** *Rhinolophus hipposideros* sampling sites for the whole genome sequencing study and measures of genomic diversity (ROH: Runs of Homozygosity; Ho: Observed Heterozygosity; Theta: genome wide heterozygosity).

| Site Code | County | Date Sampled | *N* | Theta (median) | Variant *Ho* (median) | ROH (bp) | Segments in ROH (%) | ALAN |
|---|---|---|---|---|---|---|---|---|
| BE | Berkshire | 15/08/2024 | 11 | 0.0024 | 0.128 | 1666670 | 0.2378 | 13.2 |
| BR | Bristol | 15/08/2024 | 12 | 0.0021 | 0.121 | 1666670 | 0.2378 | 253.6 |
| CO1 | Cornwall | 18/09/2024 | 12 | 0.0021 | 0.108 | 1666670 | 0.2378 | 20.3 |
| CO2 | Cornwall | 18/09/2024 | 12 | 0.0021 | 0.121 | 1583335 | 0.2378 | 28.2 |
| CO3 | Cornwall | 17/09/2024 | 8 | 0.0021 | 0.125 | 1666670 | 0.2378 | 15.9 |
| DE1 | Devon | 27/09/2024 | 9 | 0.0022 | 0.120 | 1666670 | 0.2378 | 11.1 |
| DE2 | Devon | 20/09/2024 | 10 | 0.0022 | 0.126 | 1416665 | 0.2141 | 11.0 |
| DE3 | Devon | 12/08/2024 | 12 | 0.0024 | 0.118 | 1666670 | 0.2378 | 16.5 |
| GL1 | Gloucestershire | 15/09/2024 | 11 | 0.0022 | 0.124 | 1666670 | 0.2378 | 15.1 |
| GL2 | Gloucestershire | 16/09/2024 | 9 | 0.0022 | 0.143 | 1666670 | 0.2378 | 14.0 |
| HE | Herefordshire | 19/09/2024 | 8 | 0.0022 | 0.133 | 1666670 | 0.2378 | 13.7 |
| NW1 | North Wales, Gwynedd | 26/09/2024 | 12 | 0.0021 | 0.115 | 1666670 | 0.2378 | 11.5 |
| NW2 | North Wales, Gwynedd | 30/09/2024 | 9 | 0.0021 | 0.107 | 1666670 | 0.2378 | 9.7 |
| SO1 | Somerset | 22/08/2024 | 10 | 0.0023 | 0.120 | 1666670 | 0.2378 | 24.7 |
| SO2 | Somerset | 25/08/2024 | 11 | 0.0023 | 0.135 | 1666670 | 0.2379 | 13.1 |
| SO3 | Somerset | 24/08/2024 | 12 | 0.0021 | 0.138 | 1666670 | 0.2378 | 117.3 |
| SO4 | Somerset | 26/08/2024 | 12 | 0.0023 | 0.157 | 1666670 | 0.2378 | 177.9 |
| SO5 | Somerset | 20/08/2024 | 12 | 0.0023 | 0.115 | 1666670 | 0.2378 | 131.9 |
| SW | South Wales, Monmouthshire | 25/09/2024 | 6 | 0.0022 | 0.134 | 1666670 | 0.2378 | 11.9 |

**Table S4.** Results of the statistical analysis relating measures of genetic diversity in *Rhinolophus hipposideros* (Ho: observed heterozygosity SNP variants; ROH length: runs of homozygosity average length in bp; ROH %: runs of homozygosity proportion of genome; Theta: genome-wide heterozygosity) with landscape variables.

| **Diversity metric** | **Landscape variable** | **rho** | ***P*-value** |
|---|---|---:|---:|
| Ho | Artificial light at night | -0.004 | 0.954 |
| Ho | Broadleaf woodland area | -0.049 | 0.493 |
| Ho | All woodland area | -0.053 | 0.455 |
| Ho | Distance to broadleaf woodland | 0.110 | 0.124 |
| Ho | Arable area | 0.112 | 0.115 |
| Ho | Urban area | -0.011 | 0.882 |
| ROH length | Artificial light at night | 0.053 | 0.460 |
| ROH length | Broadleaf woodland area | 0.001 | 0.990 |
| ROH length | All woodland area | -0.012 | 0.871 |
| ROH length | Distance to broadleaf woodland | -0.004 | 0.961 |
| ROH length | Arable area | -0.053 | 0.455 |
| ROH length | Urban area | 0.072 | 0.310 |
| ROH % | Artificial light at night | 0.027 | 0.707 |
| ROH % | Broadleaf woodland area | -0.041 | 0.568 |
| ROH % | All woodland area | -0.021 | 0.769 |
| ROH % | Distance to broadleaf woodland | -0.024 | 0.737 |
| ROH % | Arable area | -0.109 | 0.125 |
| ROH % | Urban area | 0.063 | 0.380 |
| Theta | Artificial light at night | -0.086 | 0.230 |
| Theta | Broadleaf woodland area | -0.215 | 0.002 |
| Theta | All woodland area | -0.215 | 0.002 |
| Theta | Distance to broadleaf woodland | 0.066 | 0.358 |
| Theta | Arable area | 0.100 | 0.162 |
| Theta | Urban area | 0.059 | 0.406 |

**Table S5.** Models compared in the landscape genetics analysis outlining the effect of landscape variables on genetic connectivity between *Rhinolophus hipposideros* colonies in Britain, including the resistance costs allocated to each landscape variable and rationale behind the cost allocation based on potential effect on movement.

| Variable | Source map | Effect on movement (rationale) | Resistance costs |
|---|---|---|---|
| Land Cover 1 | CEH 2023 Land Cover map (https://www.ceh.ac.uk/data/ukceh-land-cover-maps) | Strong effect of land cover on gene flow, strongest effect urban, only forests and grasslands do not form strong barriers. Broadleaf woodland facilitates gene flow more than conifer woodland. | Sea = 200<br>Broadleaf woodland = 1<br>Coniferous woodland = 10<br>Arable = 40<br>Improved grassland = 30<br>Semi-natural grass = 20<br>Mountain, heath = 50<br>Saltwater = 50<br>Freshwater = 20<br>Coastal = 50<br>Urban = 100 |
| Land Cover 2 | CEH 2023 Land Cover map (https://www.ceh.ac.uk/data/ukceh-land-cover-maps) | Medium effect of land cover, with urban forming the strongest barrier and no difference between forest types | Sea = 200<br>Broadleaf woodland = 1<br>Coniferous woodland = 1<br>Arable = 30<br>Improved grassland = 30<br>Semi-natural grass = 30<br>Mountain, heath = 30<br>Saltwater = 30<br>Freshwater = 20<br>Coastal = 30<br>Urban = 50 |
| Land Cover 3 | CEH 2023 Land Cover map (https://www.ceh.ac.uk/data/ukceh-land-cover-maps) | Weak effect of land cover, with urban forming stronger barrier and no difference between forest types | Sea = 200<br>Broadleaf woodland = 1<br>Coniferous woodland = 1<br>Arable = 10<br>Improved grassland = 10<br>Semi-natural grass = 10<br>Mountain, heath = 10<br>Saltwater = 10<br>Freshwater = 10<br>Coastal = 10<br>Urban = 30 |
| Null model | CEH 2023 Land Cover map (https://www.ceh.ac.uk/data/ukceh-land-cover-maps) | No effect of landscape on gene flow | Sea = 200<br>Broadleaf woodland = 1<br>Coniferous woodland = 1<br>Arable = 1<br>Improved grassland = 1<br>Semi-natural grass = 1<br>Mountain, heath = 1<br>Saltwater = 1<br>Freshwater = 1<br>Coastal = 1<br>Urban = 1 |
| Distance to broadleaf woodland | National Forest Inventory 2023 woodland map (https://www.forestresearch.gov.uk/tools-and-resources/national-forest-inventory) | The greater the distance from woodland the less likely the bats are to fly through there | 1-100 (resistance increases with distance from woodlands) |

| | | | |
|---|---|---|---|
| ALAN | NASA Black Marble (https://blackmarble.gsfc.nasa.gov) | Bats tend to avoid lit areas, therefore resistance increasing with increasing light intensity | 1-100 (resistance increases with increasing light intensity) |

**Table S6.** Results of the Maximum-Likelihood Population Effect models relating measures of genetic differentiation ($F_{ST}$, adjusted for effect of isolation by distance) and cumulative costs of crossing the different landscape variables between *Rhinolophus hipposideros* colonies in Britain (AICc: Akaike Information Criterion corrected for small sample sizes; BIC: Bayesian Information Criterion). The best supported model is highlighted in bold.

| **Model design** | **Conditional $R^2$** | **Marginal $R^2$** | **AICc weights** | **BIC weights** |
|---|---|---|---|---|
| **$F_{ST}$/km ~ Distance to broadleaf woodland + ALAN + (1\|Site1)** | **0.758** | **0.330** | **0.738** | **0.907** |
| $F_{ST}$ /km ~ Distance to broadleaf woodland + ALAN + Land cover 1 + (1\|Site1) | 0.760 | 0.331 | 0.256 | 0.070 |
| $F_{ST}$ /km ~ Distance to broadleaf woodland + Land cover 1 + (1\|Site1) | 0.739 | 0.311 | 0.003 | 0.003 |
| $F_{ST}$ /km ~ Distance to broadleaf woodland + (1\|Site1) | 0.729 | 0.309 | 0.004 | 0.020 |
| $F_{ST}$ /km ~ ALAN + Land cover 1 + (1\|Site1) | 0.672 | 0.273 | 0.000 | 0.000 |
| $F_{ST}$ /km ~ Land cover 1 + (1\|Site1) | 0.677 | 0.276 | 0.000 | 0.000 |
| $F_{ST}$ /km ~ ALAN + (1\|Site1) | 0.476 | 0.023 | 0.000 | 0.000 |

**Table S7.** List of genes containing SNPs identified as associated with urbanisation and artificial lighting in the genotype-environmental association (GEA) analysis.

| Gene Containing Outlier SNP | Gene Containing Outlier SNP | Gene Containing Outlier SNP | Gene Containing Outlier SNP | Gene Containing Outlier SNP |
|---|---|---|---|---|
| ABI1 | CUEDC1 | HSF4 | PCDHGB7 | TENT4B |
| ACBD3 | CYP2B6 | HSPA4 | PCOLCE2 | TG |
| ADAMTS15 | CYP2S1 | IFTAP | PCSK2 | TIMP2 |
| ADAMTS9 | CYP3A4 | IGSF3 | PDE3B | TMEM129 |
| ADCY5 | CYP3A43 | IL1RAPL1 | PDE4D | TMEM132E |
| ADGRE3 | CYP3A5 | IL1RAPL2 | PDGFD | TMEM163 |
| AFF3 | CYP3A7 | INPP5D | PDILT | TMEM232 |
| AGAP1 | CYP3A7-CYP3A51P | JHY | PDLIM2 | TMEM94 |
| AGBL4 | DAW1 | JPH3 | PER3 | TNR |
| AHI1 | DCAF13 | KALRN | PGBD5 | TNS4 |
| ALG11 | DCDC2C | KAZN | PID1 | TOM1L2 |
| ALKBH8 | DCHS2 | KCND2 | PIEZO2 | TOX2 |
| ANKFN1 | DCLK2 | KCND3 | PIK3C2G | TP53RK |
| ANKRD13C | DCTN6 | KCNH7 | PIK3R6 | TRAPPC12 |
| ANKRD44 | DGKI | KCNIP4 | PKNOX2 | TRIM44 |
| ANKS1B | DHRSX | KCNMA1 | PLPP4 | TRIM69 |
| ANOS1 | DIAPH2 | KHSRP | POU6F2 | TRIT1 |
| APPBP2 | DIPK1A | KIF6 | PPA1 | TRUB2 |
| ARHGAP26 | DIPK1C | KIRREL3 | PPM1H | TSHZ3 |
| ARHGEF10 | DIPK2B | KMT5B | PPP1R36 | TSPEAR |
| ARHGEF17 | DKK2 | KRTAP4-5 | PPP1R7 | TTC28 |
| ARHGEF3 | DLG1 | KSR2 | PPP3CC | TTC6 |
| ARHGEF37 | DLG2 | LANCL2 | PPP6R3 | TTC9 |
| ARL13B | DLG5 | LBR | PRIM2 | TUBGCP4 |
| ARL8B | DLGAP3 | LINC00514 | PRKAR2A | UBR1 |
| ASIC2 | DMXL1 | LNPEP | PRKCB | ULBP1 |
| ASIC4 | DNAH17 | LRP1B | PRKCQ | USH2A |
| ASMT | DNAH8 | LRRTM4 | PRKN | USP2 |
| ASMTL | DOCK4 | LUZP2 | PROKR1 | USP28 |
| ASTN2 | DOCK5 | LYST | PRRX2 | USP46 |
| ASXL1 | DOK6 | MALRD1 | PTCD1 | UTRN |
| ATG16L2 | DOP1B | MAML3 | PTPN7 | VSNL1 |
| ATP11C | DYNC2LI1 | MANBA | PTPRG | VWCE |
| ATP23 | EDA | MAPK8IP3 | PTPRM | WAPL |
| ATP5MF | EEF1E1 | MASTL | PTPRT | WDFY4 |
| ATP6V0A4 | EGLN3 | MATN1 | PUM1 | WDR41 |
| ATP8A1 | ELMO1 | MCC | RAB18 | WDR49 |
| ATRNL1 | ENO4 | MCTP1 | RAB22A | WRN |
| AUH | EPHA3 | MDGA2 | RBFOX1 | WWOX |
| B3GAT2 | EPHA6 | MFSD14B | RBMS3 | WWTR1 |
| B4GALT5 | EPHB1 | MIB1 | RERE | XPO1 |
| BLOC1S5 | EPS15 | MID2 | RFX1 | XPO7 |
| BNC1 | ERBB4 | MLLT3 | RFX6 | ZDHHC21 |
| BTBD9 | ERC1 | MMP16 | RNF169 | ZFYVE9 |
| C12orf54 | EVC2 | MPZ | RNF220 | ZMYND11 |
| C1orf112 | EVI5 | MTFR1 | ROBO1 | ZNF169 |

| | | | | |
|---|---|---|---|---|
| C1orf158 | EVL | MTM1 | ROBO2 | ZNF423 |
| C1R | EXOC4 | MTMR6 | ROCK2 | ZNF510 |
| C4A | EYS | MUC16 | RORC | ZNF521 |
| C4B | FAM118A | MUC22 | RP1 | ZNF658 |
| CACNA2D1 | FAM174B | MYH13 | RPS6KA2 | ZNF75A |
| CACNA2D2 | FAM178B | MYLK4 | RYR2 | ZNF75D |
| CACNB4 | FAM183A | MYO18B | SAMSN1 | ZNF804B |
| CALN1 | FAM189A1 | MYO5B | SBF2 | ZNF875 |
| CAMKMT | FAM189A2 | MYO7A | SCN11A | ZNF99 |
| CAMTA1 | FAM214A | NAA20 | SCNN1B | ZW10 |
| CAPN1 | FAT3 | NAALADL2 | SCUBE1 | CTNNA3 |
| CARMIL1 | FAT4 | NANP | SDAD1 | HS6ST3 |
| CASQ2 | FBXO42 | NAV2 | SERPINB1 | PCDHGB6 |
| CCDC12 | FGD2 | NCF1 | SEZ6 | TENM2 |
| CCDC138 | FMNL3 | NFS1 | SFXN5 | |
| CCDC157 | FOXN3 | NHS | SGCZ | |
| CCDC3 | FRMD4A | NLGN1 | SGK1 | |
| CCDC60 | FSIP2 | NRG2 | SH3RF3 | |
| CDCP1 | FSTL5 | NRG3 | SHISA9 | |
| CDH12 | FYB1 | NTN3 | SLC22A8 | |
| CDH23 | GABBR2 | NUB1 | SLC24A3 | |
| CFAP47 | GALNT17 | NUP153 | SLC25A16 | |
| CFAP58 | GARRE1 | NXPE3 | SLC25A20 | |
| CFAP74 | GBE1 | OSBP2 | SLC2A1 | |
| CHD1L | GFRA2 | OSBPL5 | SLC3A2 | |
| CHD9 | GLS | OTUD7A | SLC8A3 | |
| CHST9 | GPATCH1 | PABIR3 | SLC9A9 | |
| CHSY3 | GPR143 | PADI1 | SMARCC1 | |
| CIT | GPR158 | PAPPA2 | SMOC1 | |
| CKMT2 | GPR89A | PASD1 | SNTG1 | |
| CLCN5 | GPR89B | PATJ | SORD | |
| CLPTM1L | GPSM2 | PAX3 | SPATA6 | |
| CLSTN2 | GRB14 | PBX1 | SPNS3 | |
| CMSS1 | GRID2 | PCCB | SPOCK1 | |
| CNTN4 | GRIK2 | PCDH15 | SRBD1 | |
| COBL | GRIK3 | PCDHGA1 | ST6GALNAC3 | |
| COG5 | GRIK4 | PCDHGA11 | ST7 | |
| COL22A1 | GRM7 | PCDHGA12 | STX8 | |
| COL24A1 | GSTO2 | PCDHGA2 | SUCO | |
| COL4A3 | GTF2I | PCDHGA3 | SV2A | |
| CORO7 | GTF2IRD2 | PCDHGA4 | SYN2 | |
| CP | GTF2IRD2B | PCDHGA5 | SYN3 | |
| CPQ | GUCY1A2 | PCDHGA6 | SYNGR1 | |
| CRAMP1 | GYG1 | PCDHGA7 | SYNPR | |
| CRNKL1 | GYG2 | PCDHGA8 | SYT11 | |
| CRTAM | HAUS6 | PCDHGA9 | SYT16 | |
| CRYL1 | HHLA1 | PCDHGB1 | TAF2 | |
| CSF2RA | HK2 | PCDHGB2 | TAMM41 | |
| CSMD1 | HLA-DQA1 | PCDHGB3 | TBCEL | |
| CSMD3 | HNF4A | PCDHGB4 | TCHH | |
| CTDSP2 | HPSE2 | PCDHGB5 | TECTA | |

**Table S8** – Gene set enrichment analysis results for genes with outlier SNPs in *Rhinolophus hipposideros* based on the GEA analysis. Gene set = GO Biological Process 2023.

| Term | Adjust P-value | Odds Ratio | Combine Score | GEA Genes in Gene Set | Fraction GEA Genes in Gene Set |
|---|---|---|---|---|---|
| ERBB4 Signaling Pathway (GO:0038130) | 0.030 | 62.16 | 557.20 | NRG3;ERBB4;NRG2 | 3/5 |
| Oxidative Demethylation (GO:0070989) | 0.012 | 41.53 | 448.78 | CYP3A7-CYP3A51P;CYP3A43;CYP3A4;CYP3A5 | 4/9 |
| Retinal Ganglion Cell Axon Guidance (GO:0031290) | 0.036 | 41.44 | 343.47 | ROBO2;PTPRM;EPHB1 | 3/6 |
| Synaptic Vesicle Clustering (GO:0097091) | 0.036 | 41.44 | 343.47 | NLGN1;SYN3;SYN2 | 3/6 |
| Synaptic Vesicle Localization (GO:0097479) | 0.036 | 41.44 | 343.47 | NLGN1;SYN3;SYN2 | 3/6 |
| Lipid Hydroxylation (GO:0002933) | 0.036 | 41.44 | 343.47 | CYP3A7-CYP3A51P;CYP3A4;CYP3A5 | 3/6 |
| Positive Regulation of Synapse Assembly (GO:0051965) | 0.030 | 8.93 | 79.77 | NLGN1;GRID2;CLSTN2;DLG5;IL1RAPL1;EPHB1 | 6/35 |
| Regulation of Trans-Synaptic Signaling (GO:0099177) | 0.030 | 7.30 | 66.46 | NLGN1;GRID2;NRG3;GRIK3;GRIK4;DLGAP3;BTBD9 | 7/47 |
| Modulation of Chemical Synaptic Transmission (GO:0050804) | 0.008 | 5.27 | 65.28 | USP46;GRID2;NLGN1;MCTP1;CLSTN2;GRIK3;GRIK4;BTBD9;SYN3;SLC8A3;GRM7;NRG3;DLGAP3 | 13/117 |
| Regulation of Synapse Assembly (GO:0051963) | 0.030 | 6.08 | 55.07 | ROBO2;GRID2;NLGN1;CLSTN2;IL1RAPL2;DLG5;IL1RAPL1;EPHB1 | 8/64 |
| Sensory Perception of Mechanical Stimulus (GO:0050954) | 0.027 | 5.18 | 50.41 | GRM7;COL4A3;PIEZO2;PCDH15;CDH23;PAX3;ATP6V0A4;MYO7A;TSPEAR;USH2A | 10/92 |
| Cell-Cell Adhesion via Plasma-Membrane Adhesion Molecules (GO:0098742) | 0.010 | 4.17 | 47.62 | KIRREL3;PTPRT;ROBO2;GRID2;NLGN1;PCDHGB4;CRTAM;PTPRM;ROBO1;MPZ;IL1RAPL1;CDH23;CDH12;CNTN4;FAT4 | 15/171 |
| Axon Guidance (GO:0007411) | 0.036 | 3.62 | 29.89 | ROBO2;EPHA6;ANOS1;TNR;PTPRM;PRKCQ;EVL;CNTN4;EPHB1;KALRN;EPHA3;NTN3 | 12/152 |

**Table S9** – Input file for the landscape genetics analysis including measures of genetic differentiation ($F_{ST}$), Euclidean distance and cumulative resistance costs of the different landscape variables (ALAN: artificial lights at night; Wood: distance to woodland; LC: land cover; Null: null model) between *Rhinolophus hipposideros* colonies in Britain.

| Site1 | Site2 | Euclidean distance | Euclidean distance log | $F_{ST}$ | ALAN | Wood | LC1 | LC2 | LC3 | Null |
|---|---|---|---|---|---|---|---|---|---|---|
| 1 | 2 | 104.446 | 4.649 | 0.069 | 38.93 | 9.329 | 49.360 | 37.174 | 17.411 | 1.617 |
| 1 | 3 | 110.682 | 4.707 | 0.053 | 10.11 | 9.949 | 36.486 | 33.497 | 13.237 | 1.625 |
| 1 | 4 | 95.382 | 4.558 | 0.030 | 47.32 | 12.369 | 48.327 | 46.436 | 19.304 | 2.150 |
| 1 | 5 | 171.473 | 5.144 | 0.129 | 10.30 | 10.833 | 34.048 | 31.400 | 14.054 | 2.136 |
| 1 | 6 | 145.927 | 4.983 | 0.055 | 9.94 | 10.025 | 36.672 | 33.497 | 13.785 | 1.837 |
| 1 | 7 | 76.232 | 4.334 | 0.020 | 6.43 | 6.693 | 27.472 | 26.541 | 10.397 | 1.266 |
| 1 | 8 | 98.735 | 4.592 | 0.042 | 18.85 | 8.593 | 30.259 | 28.399 | 12.202 | 1.601 |
| 1 | 9 | 208.601 | 5.340 | 0.108 | 11.10 | 12.253 | 39.731 | 35.265 | 15.675 | 2.333 |
| 1 | 10 | 97.304 | 4.578 | 0.034 | 27.29 | 8.618 | 40.133 | 34.941 | 14.406 | 1.493 |
| 1 | 11 | 187.025 | 5.231 | 0.053 | 10.52 | 10.355 | 39.296 | 36.650 | 14.719 | 1.877 |
| 1 | 12 | 93.394 | 4.537 | 0.048 | 7.55 | 7.397 | 29.643 | 28.406 | 11.143 | 1.314 |
| 1 | 13 | 64.880 | 4.173 | 0.027 | 4.92 | 4.764 | 20.064 | 19.111 | 7.379 | 0.918 |
| 1 | 14 | 137.782 | 4.926 | 0.075 | 13.38 | 12.623 | 47.800 | 44.672 | 18.170 | 2.113 |
| 1 | 15 | 116.324 | 4.756 | 0.044 | 9.79 | 9.451 | 33.850 | 32.571 | 13.554 | 1.845 |
| 1 | 16 | 172.232 | 5.149 | 0.051 | 10.22 | 9.837 | 38.756 | 36.166 | 14.476 | 1.841 |
| 1 | 17 | 49.512 | 3.902 | 0.019 | 6.24 | 6.871 | 31.338 | 29.568 | 10.770 | 1.160 |
| 1 | 18 | 84.505 | 4.437 | 0.044 | 7.86 | 8.936 | 31.554 | 31.466 | 12.138 | 1.456 |
| 1 | 19 | 73.239 | 4.294 | 0.048 | 22.35 | 11.729 | 36.979 | 33.856 | 13.405 | 1.526 |
| 2 | 3 | 11.567 | 2.448 | 0.053 | 35.46 | 4.744 | 35.430 | 22.409 | 10.540 | 0.597 |
| 2 | 4 | 140.831 | 4.948 | 0.077 | 81.30 | 16.366 | 70.703 | 57.013 | 27.262 | 2.722 |
| 2 | 5 | 165.318 | 5.108 | 0.137 | 37.74 | 7.727 | 43.531 | 29.312 | 13.925 | 1.181 |
| 2 | 6 | 72.027 | 4.277 | 0.062 | 36.73 | 6.190 | 41.369 | 27.251 | 12.542 | 0.853 |
| 2 | 7 | 72.274 | 4.280 | 0.062 | 36.39 | 6.777 | 38.874 | 26.979 | 12.975 | 0.973 |
| 2 | 8 | 5.978 | 1.788 | 0.029 | 43.21 | 3.121 | 28.220 | 16.018 | 9.104 | 0.532 |
| 2 | 9 | 204.099 | 5.319 | 0.118 | 38.52 | 9.121 | 49.216 | 33.208 | 15.521 | 1.375 |

| | | | | | | | | | | |
|---|---|---|---|---|---|---|---|---|---|---|
| 2 | 10 | 56.745 | 4.039 | 0.051 | 54.99 | 6.018 | 45.491 | 29.643 | 14.237 | 0.812 |
| 2 | 11 | 91.151 | 4.513 | 0.058 | 37.08 | 6.454 | 43.402 | 30.013 | 13.410 | 0.915 |
| 2 | 12 | 186.749 | 5.230 | 0.095 | 42.09 | 12.307 | 54.655 | 41.500 | 20.218 | 2.084 |
| 2 | 13 | 122.840 | 4.811 | 0.072 | 38.98 | 8.795 | 43.412 | 30.443 | 15.424 | 1.478 |
| 2 | 14 | 230.915 | 5.442 | 0.117 | 47.764 | 17.359 | 72.041 | 57.042 | 26.996 | 2.877 |
| 2 | 15 | 55.568 | 4.018 | 0.055 | 36.743 | 5.793 | 39.445 | 27.083 | 12.527 | 0.861 |
| 2 | 16 | 70.508 | 4.256 | 0.056 | 36.732 | 5.876 | 42.480 | 29.194 | 13.092 | 0.882 |
| 2 | 17 | 152.132 | 5.025 | 0.081 | 40.910 | 12.064 | 57.299 | 43.618 | 20.201 | 1.965 |
| 2 | 18 | 29.463 | 3.383 | 0.044 | 35.075 | 5.588 | 35.630 | 24.939 | 11.570 | 0.743 |
| 2 | 19 | 31.459 | 3.449 | 0.042 | 49.975 | 8.555 | 42.929 | 29.321 | 13.661 | 0.917 |
| 3 | 4 | 138.761 | 4.933 | 0.061 | 52.473 | 16.980 | 57.755 | 53.258 | 23.076 | 2.729 |
| 3 | 5 | 176.508 | 5.173 | 0.117 | 8.959 | 8.378 | 30.662 | 25.711 | 9.777 | 1.200 |
| 3 | 6 | 79.927 | 4.381 | 0.039 | 7.929 | 6.829 | 28.386 | 23.599 | 8.361 | 0.867 |
| 3 | 7 | 69.798 | 4.246 | 0.045 | 7.538 | 7.366 | 25.844 | 23.101 | 8.743 | 0.976 |
| 3 | 8 | 16.438 | 2.800 | 0.024 | 16.481 | 4.631 | 19.251 | 15.848 | 6.279 | 0.662 |
| 3 | 9 | 215.145 | 5.371 | 0.098 | 9.735 | 9.770 | 36.333 | 29.597 | 11.368 | 1.394 |
| 3 | 10 | 50.660 | 3.925 | 0.035 | 26.003 | 6.489 | 31.829 | 25.151 | 9.792 | 0.796 |
| 3 | 11 | 92.707 | 4.529 | 0.037 | 8.068 | 6.951 | 29.675 | 25.650 | 8.980 | 0.905 |
| 3 | 12 | 189.454 | 5.244 | 0.078 | 13.265 | 12.927 | 41.766 | 37.810 | 16.043 | 2.092 |
| 3 | 13 | 123.003 | 4.812 | 0.057 | 10.160 | 9.411 | 30.501 | 26.720 | 11.243 | 1.486 |
| 3 | 14 | 233.145 | 5.452 | 0.100 | 18.944 | 17.980 | 59.155 | 53.359 | 22.822 | 2.885 |
| 3 | 15 | 66.029 | 4.190 | 0.030 | 7.998 | 6.485 | 26.773 | 23.682 | 8.453 | 0.890 |
| 3 | 16 | 69.571 | 4.242 | 0.034 | 7.690 | 6.339 | 28.580 | 24.661 | 8.604 | 0.867 |
| 3 | 17 | 157.181 | 5.057 | 0.063 | 12.090 | 12.684 | 44.413 | 39.932 | 16.026 | 1.973 |
| 3 | 18 | 29.500 | 3.384 | 0.028 | 6.416 | 6.370 | 23.178 | 21.578 | 7.525 | 0.762 |
| 3 | 19 | 37.663 | 3.629 | 0.027 | 21.358 | 9.453 | 30.711 | 26.188 | 9.689 | 0.945 |
| 4 | 5 | 262.457 | 5.570 | 0.133 | 52.822 | 18.051 | 56.774 | 52.419 | 24.176 | 3.247 |
| 4 | 6 | 207.537 | 5.335 | 0.063 | 52.407 | 17.158 | 58.626 | 53.847 | 23.763 | 2.944 |
| 4 | 7 | 68.963 | 4.234 | 0.023 | 48.700 | 13.509 | 47.557 | 45.144 | 19.954 | 2.359 |

| | | | | | | | | | | |
|---|---|---|---|---|---|---|---|---|---|---|
| 4 | 8 | 137.880 | 4.926 | 0.049 | 61.208 | 15.623 | 51.615 | 48.252 | 22.053 | 2.705 |
| 4 | 9 | 300.927 | 5.707 | 0.115 | 53.620 | 19.466 | 62.434 | 56.270 | 25.791 | 3.443 |
| 4 | 10 | 92.071 | 4.523 | 0.040 | 69.602 | 15.550 | 60.784 | 54.134 | 24.120 | 2.594 |
| 4 | 11 | 231.375 | 5.444 | 0.060 | 52.937 | 17.450 | 60.968 | 56.758 | 24.644 | 2.984 |
| 4 | 12 | 82.600 | 4.414 | 0.029 | 49.679 | 14.680 | 50.743 | 48.118 | 21.028 | 2.474 |
| 4 | 13 | 30.507 | 3.418 | 0.005 | 45.740 | 10.287 | 35.799 | 33.788 | 15.201 | 1.786 |
| 4 | 14 | 114.025 | 4.736 | 0.054 | 55.526 | 19.915 | 69.014 | 64.475 | 28.067 | 3.275 |
| 4 | 15 | 181.800 | 5.203 | 0.054 | 52.275 | 16.608 | 55.963 | 53.056 | 23.564 | 2.953 |
| 4 | 16 | 207.893 | 5.337 | 0.059 | 52.638 | 16.918 | 60.331 | 56.186 | 24.382 | 2.947 |
| 4 | 17 | 92.543 | 4.528 | 0.035 | 48.580 | 14.576 | 53.991 | 50.789 | 21.178 | 2.377 |
| 4 | 18 | 111.381 | 4.713 | 0.053 | 50.200 | 15.910 | 52.545 | 50.970 | 21.905 | 2.559 |
| 4 | 19 | 117.071 | 4.763 | 0.055 | 64.712 | 18.725 | 58.153 | 53.496 | 23.185 | 2.629 |
| 5 | 6 | 114.098 | 4.737 | 0.107 | 5.454 | 6.530 | 27.118 | 22.486 | 8.306 | 1.037 |
| 5 | 7 | 212.681 | 5.360 | 0.119 | 8.658 | 9.011 | 28.253 | 25.113 | 10.732 | 1.518 |
| 5 | 8 | 162.270 | 5.089 | 0.111 | 18.291 | 7.173 | 24.665 | 20.691 | 8.851 | 1.199 |
| 5 | 9 | 39.645 | 3.680 | 0.046 | 3.672 | 5.469 | 19.964 | 15.953 | 6.087 | 0.843 |
| 5 | 10 | 212.929 | 5.361 | 0.117 | 27.626 | 8.647 | 37.011 | 29.672 | 12.458 | 1.356 |
| 5 | 11 | 160.229 | 5.077 | 0.102 | 7.376 | 7.401 | 30.586 | 26.342 | 9.759 | 1.173 |
| 5 | 12 | 256.566 | 5.547 | 0.147 | 13.354 | 13.737 | 38.852 | 35.367 | 16.743 | 2.600 |
| 5 | 13 | 233.094 | 5.451 | 0.129 | 10.500 | 10.480 | 29.259 | 25.699 | 12.336 | 2.004 |
| 5 | 14 | 296.491 | 5.692 | 0.171 | 18.915 | 18.704 | 55.755 | 50.522 | 23.399 | 3.389 |
| 5 | 15 | 111.725 | 4.716 | 0.100 | 5.023 | 5.780 | 24.389 | 21.610 | 7.954 | 0.997 |
| 5 | 16 | 173.033 | 5.153 | 0.102 | 7.338 | 7.147 | 30.621 | 26.393 | 9.796 | 1.190 |
| 5 | 17 | 206.901 | 5.332 | 0.142 | 12.204 | 13.512 | 41.532 | 37.520 | 16.750 | 2.482 |
| 5 | 18 | 180.318 | 5.195 | 0.114 | 8.177 | 8.675 | 28.082 | 25.832 | 10.108 | 1.320 |
| 5 | 19 | 163.515 | 5.097 | 0.115 | 22.725 | 11.408 | 33.719 | 28.626 | 11.782 | 1.481 |
| 6 | 7 | 141.425 | 4.952 | 0.048 | 7.986 | 7.867 | 28.339 | 25.062 | 9.879 | 1.203 |
| 6 | 8 | 72.636 | 4.285 | 0.033 | 17.382 | 5.686 | 22.777 | 18.813 | 7.546 | 0.877 |
| 6 | 9 | 148.452 | 5.000 | 0.086 | 6.215 | 7.911 | 32.757 | 26.355 | 9.882 | 1.228 |

| | | | | | | | | | | |
|---|---|---|---|---|---|---|---|---|---|---|
| 6 | 10 | 128.730 | 4.858 | 0.040 | 26.769 | 7.280 | 35.843 | 28.501 | 11.303 | 1.033 |
| 6 | 11 | 53.174 | 3.974 | 0.022 | 6.182 | 5.799 | 27.706 | 23.865 | 8.252 | 0.845 |
| 6 | 12 | 238.268 | 5.473 | 0.079 | 13.036 | 12.973 | 41.852 | 37.737 | 16.550 | 2.303 |
| 6 | 13 | 185.039 | 5.221 | 0.059 | 10.089 | 9.589 | 31.246 | 27.221 | 11.926 | 1.701 |
| 6 | 14 | 283.032 | 5.646 | 0.102 | 18.642 | 17.985 | 59.068 | 53.140 | 23.277 | 3.094 |
| 6 | 15 | 29.676 | 3.390 | 0.010 | 4.384 | 4.542 | 22.698 | 19.942 | 6.895 | 0.736 |
| 6 | 16 | 59.298 | 4.083 | 0.019 | 6.137 | 5.508 | 27.664 | 23.822 | 8.244 | 0.853 |
| 6 | 17 | 195.016 | 5.273 | 0.064 | 11.876 | 12.737 | 44.495 | 39.857 | 16.541 | 2.184 |
| 6 | 18 | 98.895 | 4.594 | 0.036 | 7.348 | 7.309 | 27.004 | 24.724 | 8.995 | 1.001 |
| 6 | 19 | 90.467 | 4.505 | 0.036 | 21.969 | 10.127 | 33.188 | 28.066 | 10.811 | 1.167 |
| 7 | 8 | 69.821 | 4.246 | 0.036 | 16.228 | 6.023 | 19.753 | 18.215 | 7.769 | 0.959 |
| 7 | 9 | 252.267 | 5.530 | 0.099 | 9.447 | 10.417 | 33.947 | 29.005 | 12.339 | 1.713 |
| 7 | 10 | 27.103 | 3.300 | 0.020 | 24.322 | 5.520 | 27.068 | 22.290 | 9.238 | 0.808 |
| 7 | 11 | 162.430 | 5.090 | 0.044 | 8.297 | 8.039 | 30.186 | 27.547 | 10.606 | 1.237 |
| 7 | 12 | 128.130 | 4.853 | 0.046 | 9.638 | 9.723 | 32.908 | 30.972 | 13.264 | 1.733 |
| 7 | 13 | 56.224 | 4.029 | 0.021 | 6.361 | 5.920 | 20.301 | 18.571 | 8.090 | 1.115 |
| 7 | 14 | 169.406 | 5.132 | 0.071 | 15.393 | 14.862 | 50.625 | 46.826 | 20.146 | 2.529 |
| 7 | 15 | 118.104 | 4.772 | 0.038 | 7.924 | 7.391 | 25.986 | 24.537 | 9.777 | 1.216 |
| 7 | 16 | 139.057 | 4.935 | 0.045 | 7.960 | 7.467 | 29.356 | 26.796 | 10.289 | 1.198 |
| 7 | 17 | 108.279 | 4.685 | 0.033 | 8.448 | 9.466 | 35.564 | 33.096 | 13.232 | 1.614 |
| 7 | 18 | 43.060 | 3.763 | 0.036 | 5.104 | 6.134 | 19.528 | 19.879 | 7.329 | 0.811 |
| 7 | 19 | 52.638 | 3.963 | 0.040 | 19.783 | 9.146 | 25.871 | 23.057 | 8.855 | 0.915 |
| 8 | 9 | 201.270 | 5.305 | 0.089 | 19.071 | 8.570 | 30.361 | 24.595 | 10.451 | 1.394 |
| 8 | 10 | 56.389 | 4.032 | 0.027 | 34.989 | 5.345 | 26.817 | 21.232 | 9.189 | 0.812 |
| 8 | 11 | 95.038 | 4.554 | 0.031 | 17.736 | 5.983 | 25.169 | 21.861 | 8.522 | 0.943 |
| 8 | 12 | 181.878 | 5.203 | 0.068 | 22.018 | 11.574 | 35.573 | 32.738 | 15.014 | 2.068 |
| 8 | 13 | 118.971 | 4.779 | 0.046 | 18.888 | 8.052 | 24.303 | 21.666 | 10.211 | 1.462 |
| 8 | 14 | 226.169 | 5.421 | 0.091 | 27.711 | 16.631 | 52.967 | 48.285 | 21.795 | 2.861 |
| 8 | 15 | 53.917 | 3.987 | 0.025 | 17.333 | 5.246 | 20.564 | 18.445 | 7.439 | 0.877 |

| | | | | | | | | | | |
|---|---|---|---|---|---|---|---|---|---|---|
| 8 | 16 | 75.348 | 4.322 | 0.028 | 17.396 | 5.417 | 24.341 | 21.119 | 8.233 | 0.912 |
| 8 | 17 | 146.638 | 4.988 | 0.053 | 20.841 | 11.331 | 38.214 | 34.855 | 14.996 | 1.949 |
| 8 | 18 | 26.764 | 3.287 | 0.015 | 14.540 | 4.603 | 15.996 | 15.782 | 6.203 | 0.717 |
| 8 | 19 | 26.050 | 3.260 | 0.013 | 29.367 | 7.437 | 23.003 | 19.953 | 8.196 | 0.881 |
| 9 | 10 | 252.497 | 5.531 | 0.097 | 28.407 | 10.043 | 42.690 | 33.558 | 14.054 | 1.550 |
| 9 | 11 | 190.002 | 5.247 | 0.080 | 8.134 | 8.761 | 36.158 | 30.151 | 11.310 | 1.359 |
| 9 | 12 | 290.529 | 5.672 | 0.129 | 14.153 | 15.150 | 44.406 | 39.114 | 18.350 | 2.796 |
| 9 | 13 | 271.270 | 5.603 | 0.110 | 11.298 | 11.896 | 34.922 | 29.554 | 13.951 | 2.200 |
| 9 | 14 | 328.832 | 5.796 | 0.155 | 19.713 | 20.114 | 61.243 | 54.211 | 24.998 | 3.586 |
| 9 | 15 | 149.677 | 5.008 | 0.075 | 5.808 | 7.197 | 30.131 | 25.555 | 9.570 | 1.197 |
| 9 | 16 | 205.992 | 5.328 | 0.082 | 8.099 | 8.514 | 36.216 | 30.222 | 11.357 | 1.378 |
| 9 | 17 | 241.030 | 5.485 | 0.120 | 13.004 | 14.929 | 47.122 | 41.302 | 18.362 | 2.678 |
| 9 | 18 | 219.844 | 5.393 | 0.094 | 8.960 | 10.074 | 33.778 | 29.733 | 11.711 | 1.515 |
| 9 | 19 | 203.132 | 5.314 | 0.095 | 23.511 | 12.813 | 39.437 | 32.544 | 13.390 | 1.676 |
| 10 | 11 | 142.241 | 4.958 | 0.037 | 26.866 | 7.289 | 36.942 | 30.298 | 11.813 | 1.055 |
| 10 | 12 | 155.228 | 5.045 | 0.058 | 30.473 | 11.616 | 45.455 | 39.285 | 17.236 | 1.961 |
| 10 | 13 | 82.975 | 4.419 | 0.037 | 27.294 | 7.983 | 33.656 | 27.693 | 12.293 | 1.350 |
| 10 | 14 | 196.278 | 5.280 | 0.082 | 36.185 | 16.705 | 62.979 | 54.954 | 24.056 | 2.755 |
| 10 | 15 | 109.554 | 4.696 | 0.034 | 26.778 | 6.899 | 33.915 | 28.350 | 11.322 | 1.054 |
| 10 | 16 | 117.171 | 4.764 | 0.037 | 26.493 | 6.667 | 35.890 | 29.332 | 11.434 | 1.013 |
| 10 | 17 | 133.932 | 4.897 | 0.046 | 29.291 | 11.368 | 48.112 | 41.412 | 17.213 | 1.842 |
| 10 | 18 | 32.796 | 3.490 | 0.028 | 24.110 | 5.710 | 26.892 | 23.176 | 8.998 | 0.717 |
| 10 | 19 | 49.663 | 3.905 | 0.031 | 39.044 | 9.108 | 34.987 | 28.253 | 11.235 | 0.900 |
| 11 | 12 | 275.533 | 5.619 | 0.079 | 13.638 | 13.311 | 44.478 | 40.887 | 17.493 | 2.343 |
| 11 | 13 | 213.951 | 5.366 | 0.057 | 10.625 | 9.883 | 33.668 | 30.195 | 12.814 | 1.740 |
| 11 | 14 | 320.159 | 5.769 | 0.102 | 19.274 | 18.336 | 61.743 | 56.331 | 24.236 | 3.135 |
| 11 | 15 | 77.801 | 4.354 | 0.013 | 6.506 | 5.768 | 27.616 | 25.184 | 8.807 | 0.924 |
| 11 | 16 | 27.948 | 3.330 | 0.006 | 4.346 | 4.185 | 21.962 | 19.844 | 6.701 | 0.677 |
| 11 | 17 | 236.481 | 5.466 | 0.064 | 12.472 | 13.073 | 47.127 | 43.013 | 17.482 | 2.225 |

| | | | | | | | | | | |
|---|---|---|---|---|---|---|---|---|---|---|
| 11 | 18 | 120.448 | 4.791 | 0.032 | 7.599 | 7.487 | 28.845 | 27.248 | 9.747 | 1.043 |
| 11 | 19 | 119.829 | 4.786 | 0.035 | 22.339 | 10.415 | 35.460 | 30.979 | 11.669 | 1.213 |
| 12 | 13 | 73.408 | 4.296 | 0.032 | 7.464 | 7.287 | 23.229 | 21.600 | 9.444 | 1.279 |
| 12 | 14 | 44.811 | 3.802 | 0.021 | 10.472 | 8.354 | 29.473 | 26.456 | 10.984 | 1.286 |
| 12 | 15 | 208.913 | 5.342 | 0.072 | 12.882 | 12.396 | 39.059 | 36.833 | 16.317 | 2.311 |
| 12 | 16 | 257.210 | 5.550 | 0.076 | 13.351 | 12.796 | 43.953 | 40.416 | 17.255 | 2.307 |
| 12 | 17 | 49.665 | 3.905 | 0.043 | 6.722 | 7.294 | 26.362 | 24.482 | 9.599 | 1.117 |
| 12 | 18 | 160.029 | 5.075 | 0.070 | 11.046 | 11.937 | 36.935 | 35.864 | 14.980 | 1.925 |
| 12 | 19 | 156.130 | 5.051 | 0.074 | 25.547 | 14.734 | 42.423 | 38.320 | 16.268 | 1.995 |
| 13 | 14 | 113.312 | 4.730 | 0.058 | 13.312 | 12.517 | 41.534 | 37.976 | 16.480 | 2.079 |
| 13 | 15 | 157.852 | 5.062 | 0.048 | 9.954 | 9.037 | 28.542 | 26.399 | 11.724 | 1.710 |
| 13 | 16 | 192.487 | 5.260 | 0.056 | 10.327 | 9.352 | 33.054 | 29.640 | 12.554 | 1.704 |
| 13 | 17 | 67.752 | 4.216 | 0.035 | 6.326 | 7.116 | 26.229 | 24.003 | 9.505 | 1.174 |
| 13 | 18 | 93.880 | 4.542 | 0.048 | 7.875 | 8.334 | 25.257 | 24.392 | 10.056 | 1.315 |
| 13 | 19 | 95.506 | 4.559 | 0.052 | 22.380 | 11.144 | 30.774 | 26.835 | 11.321 | 1.385 |
| 14 | 15 | 253.635 | 5.536 | 0.097 | 18.480 | 17.401 | 56.258 | 52.221 | 23.037 | 3.102 |
| 14 | 16 | 301.422 | 5.709 | 0.100 | 18.993 | 17.827 | 61.242 | 55.882 | 24.004 | 3.099 |
| 14 | 17 | 91.429 | 4.516 | 0.070 | 12.582 | 12.539 | 45.004 | 41.160 | 16.694 | 1.912 |
| 14 | 18 | 203.651 | 5.316 | 0.095 | 16.758 | 17.019 | 54.434 | 51.509 | 21.797 | 2.719 |
| 14 | 19 | 200.566 | 5.301 | 0.098 | 31.260 | 19.814 | 59.924 | 53.977 | 23.094 | 2.790 |
| 15 | 16 | 74.791 | 4.315 | 0.013 | 6.431 | 5.436 | 27.357 | 24.962 | 8.739 | 0.926 |
| 15 | 17 | 165.350 | 5.108 | 0.056 | 11.724 | 12.162 | 41.693 | 38.947 | 16.309 | 2.192 |
| 15 | 18 | 77.517 | 4.351 | 0.028 | 7.311 | 6.873 | 24.791 | 24.326 | 8.918 | 1.012 |
| 15 | 19 | 65.633 | 4.184 | 0.027 | 21.879 | 9.616 | 30.651 | 27.383 | 10.655 | 1.172 |
| 16 | 17 | 221.084 | 5.399 | 0.060 | 12.184 | 12.558 | 46.603 | 42.541 | 17.243 | 2.189 |
| 16 | 18 | 98.648 | 4.592 | 0.031 | 7.248 | 6.902 | 27.948 | 26.440 | 9.420 | 1.006 |
| 16 | 19 | 101.266 | 4.618 | 0.031 | 22.007 | 9.860 | 34.686 | 30.291 | 11.372 | 1.178 |
| 17 | 18 | 129.132 | 4.861 | 0.055 | 9.864 | 11.688 | 39.578 | 37.979 | 14.955 | 1.806 |
| 17 | 19 | 120.673 | 4.793 | 0.057 | 24.37 | 14.485 | 45.056 | 40.43 | 16.240 | 1.876 |
| 18 | 19 | 18.445 | 2.915 | 0.016 | 18.26 | 7.858 | 23.370 | 21.98 | 7.850 | 0.741 |

#### SI References

1. S. Chen, Ultrafast one-pass FASTQ data preprocessing, quality control, and deduplication using fastp. *iMeta* **2**, e107 (2023).
2. N. L. Bray, H. Pimentel, P. Melsted, L. Pachter, Near-optimal probabilistic RNA-seq quantification. *Nat Biotechnol* **34**, 525–527 (2016).
3. B. Langmead, S. L. Salzberg, Fast gapped-read alignment with Bowtie 2. *Nat Methods* **9**, 357–359 (2012).
4. P. Danecek, *et al.*, Twelve years of SAMtools and BCFtools. *Gigascience* **10**, giab008 (2021).
5. E. Garrison, G. Marth, Haplotype-based variant detection from short-read sequencing. [Preprint] (2012). Available at: http://arxiv.org/abs/1207.3907 [Accessed 22 December 2025].
